# Ventral tegmental area astrocytes orchestrate avoidance and approach behavior

**DOI:** 10.1101/401919

**Authors:** J. A. Gomez, J. M. Perkins, G. M. Beaudoin, N. B. Cook, S. A. Quraishi, Erin A Szoeke, K. Thangamani, C. W. Tschumi, M. J. Wanat, A. M. Maroof, M. J. Beckstead, P. A. Rosenberg, C. A. Paladini

**Affiliations:** University of Texas at San Antonio Neurosciences Institute, Department of Biology, University of Texas at San Antonio, San Antonio, TX 78249; Trinity University, Department of Biology, San Antonio, TX 78212.; Aging & Metabolism Research Program, Oklahoma Medical Research Foundation, Oklahoma City, OK 73104; Department of Neurology and the F.M. Kirby Neurobiology Center, Boston Children’s Hospital, Boston, MA, 02115

## Abstract

The ventral tegmental area (VTA) is a heterogeneous midbrain structure, containing neurons and astrocytes, that coordinates approach and avoidance behaviors by integrating activity from numerous afferents. Within neuron-astrocyte networks, astrocytes control signals from distinct afferents in a circuit-specific manner, but whether this capacity scales up to drive motivated behavior has been undetermined. Using genetic and optical dissection strategies in vitro and during behavior we report that VTA astrocytes tune glutamatergic signaling selectively on local inhibitory neurons to drive a functional circuit for learned avoidance. In this circuit, VTA astrocytes facilitate excitation of local GABA neurons to increase inhibition of dopamine neurons. The increased inhibition of dopamine neurons elicits real-time and learned avoidance behavior that is sufficient to impede expression of learned preference for reward. Despite the large number of functions performed by astrocytes, loss of one glutamate transporter (GLT-1) from VTA astrocytes selectively blocks these avoidance behaviors and spares preference for reward. Thus, VTA astrocytes selectively regulate excitation of local GABA neurons to drive a distinct learned avoidance circuit that opposes learned approach behavior.

## MAIN

The ability to coordinate approach and avoidance actions in different environments can determine individual survival. Avoidance learning is an adaptive mechanism, but can become maladaptive and lead to the development and persistence of many specific anxiety disorders like post-traumatic stress and social anxiety disorders ^1^. Dopamine signaling from the ventral tegmental area (VTA) mediates approach and avoidance learning ^2^. However, the VTA is a heterogeneous structure containing astrocytes and neurons that receive mixed and distributed signals from various independent afferents conveying information for cues that drive both approach and avoidance ^3^,^4^. For example, both aversive stimuli and reward expectation increase VTA GABA neuron firing and can inhibit dopamine neurons, leading to avoidance and reduction in approach behaviors ^5–9^. A mechanism capable of separating different signals underlying reward and aversion within the VTA is therefore necessary to appropriately coordinate approach and avoidance, respectively. By selectively affecting activity of different neurons, VTA astrocytes may be positioned to coordinate avoidance and approach behaviors.

Astrocytes control glutamate availability at synapses of actively firing afferents primarily by the glutamate transporter, GLT-1 (Eaat2, Slc1a2) ^10^. The contribution of glutamate transporters to signaling in neuron-astrocyte networks varies considerably among synapses and often changes depending on afferent activity, local ionic environment, and structure of the synapse ^10–14^. Thus, rather than indiscriminately clearing glutamate, astrocytes are diverse regulators of neuronal processing that can delineate synapse-specific control to circuits in response to neuronal afferent activity ^13–20^. Astrocytes express receptors and transporters that are activated by synaptically released neurotransmitters from neuronal afferents ^14^,^21–24^. In response to afferent activity, astrocytes display a phasic increase in cytoplasmic Na^+^ and H^+^, and a decrease in K^+^ concentrations, among other ions ^12^,^25–28^. The ensuing fast changes in electrochemical gradient generate biophysical fluctuations that slow down astrocytic clearance of extracellular glutamate ^14^,^28–30^. Thus, astrocytes are ideally suited to dynamically control glutamate at specific synapses throughout a wide concentration range ^31–33^, thereby affecting postsynaptic neuronal activity and potentially driving behavior.

In the present study, we examined the role of astrocytes in the VTA circuit and behavior. We found that VTA astrocytes 1) selectively facilitate glutamatergic excitation of local GABA neurons through a mechanism that requires astrocytic GLT-1; 2) generate a subsequent GABA neuron-dependent inhibition of dopamine neurons; and 3) consequently drive learned avoidance that is sufficient to block conditioned place preference for cocaine. Conditional mutants lacking GLT-1 in VTA astrocytes fail to express avoidance behaviors, but maintain preference for reward. These experiments establish that VTA astrocytes selectively leverage glutamatergic excitation of local GABA neurons to dynamically define a privileged neuron-astrocyte circuit underlying behaviors associated with aversion, but not reward.

## RESULTS

### Astrocytes modulate excitation of VTA GABA neurons and inhibition of dopamine neurons

To investigate the mechanism by which astrocytes regulate the circuitry of the VTA we obtained ex vivo recordings of VTA astrocytes (Fig. 1a). We first determined that neuronal afferent stimulation elicits action potential-dependent currents in VTA astrocytes (Fig. 1, S1). Electrical stimulation of the VTA during wholecell voltage-clamp recordings of astrocytes resulted in a lidocaine-sensitive inward current (Fig. 1 b-c). A significant portion of the inward current (35% ± 11%) was sensitive to the glutamate transporter blocker, TFB-TBOA (15 μM), indicating that, similar to other reports ^14^, neuronal afferent input activates glutamate receptors and glutamate transporter activity on VTA astrocytes (Fig. 1 d-e, S1).

**Figure 1.**
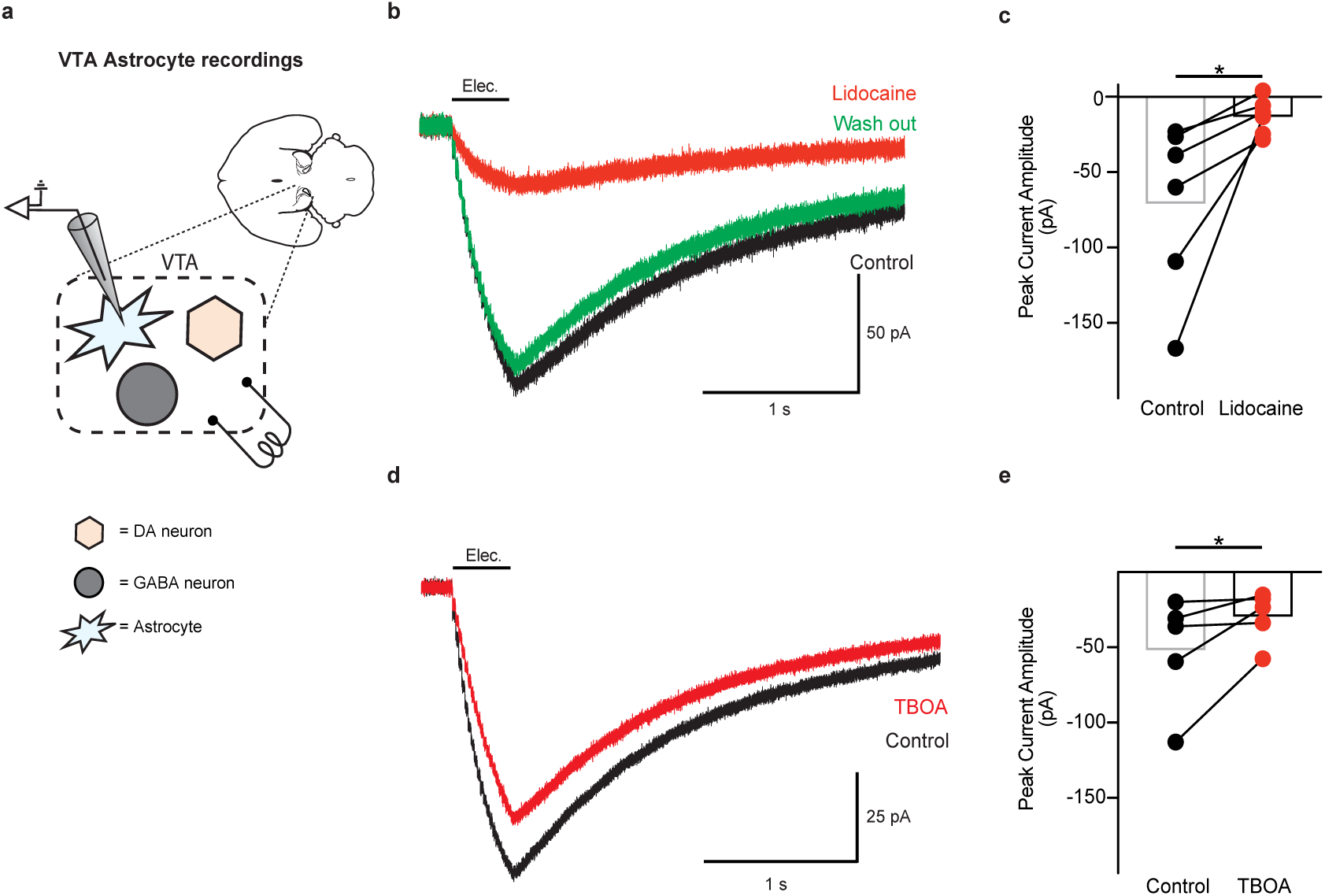
Neuronal afferent activity changes glutamate transport in VTA astrocytes. **(a)** Schematic showing the microcircuitry of the VTA and recordings performed. **(b)** Recording from a VTA astrocyte showing the current generated by electrical stimulation (black; 50 Hz, 20 pulses), the current remaining after blocking action potentials with 250 μΜ lidocaine (red), and after washing out lidocaine (green). (**c**) Summary of the currents before (black) and after lidocaine application (red; paired t-test t_(5)_ = 2.679, P = 0.04, n = 6 cells, 4 mice). (**d)** Current generated in a VTA astrocyte by electrical stimulation of afferents before (black) and after adding 15 μm TFB-TBOA (red). (**e)** Summary of the currents generated by electrical stimulation in VTA astrocytes before (black) and after (red) addition of TFB-TBOA (paired t-test t_(4)_ = 2.159, P = 0.04, n = 5 cells, 5 mice). *: P < 0.05

To mimic the activity of neuronal inputs on astrocytic glutamate transport we used an optogenetic strategy to selectively activate VTA astrocytes ^34^. Similar to afferent activation of receptors on astrocytes ^14^,^25^,^30^ (Fig. S1), photoactivation of the light-sensitive opsin, channelrhodopsin (ChR2), results in a phasic increase in cytoplasmic Na^+^ and H^+^, and a decrease in K^+^ concentrations ^35^,^36^. Therefore, owing to its cation permeability and time resolution ^35^, ChR2 presents as an ideal tool to transiently influence astrocytic GLT-1 dynamics with a temporal fidelity capable of mimicking native astrocyte gradient dynamics 28,29,37. chR2 was expressed selectively in VTA astrocytes by injecting adeno-associated virus (AAV) to drive expression of ChR2 under the gfaABCID promoter (gfaABC1D::ChR2^VTA^; Fig. 2a-b, S2) ^38^. We then confirmed that both neuronal afferent stimulation and optogenetic photoactivation of astrocytes similarly affect glutamate transporter-mediated currents in VTA astrocytes (Fig. S3).

**Figure 2.**
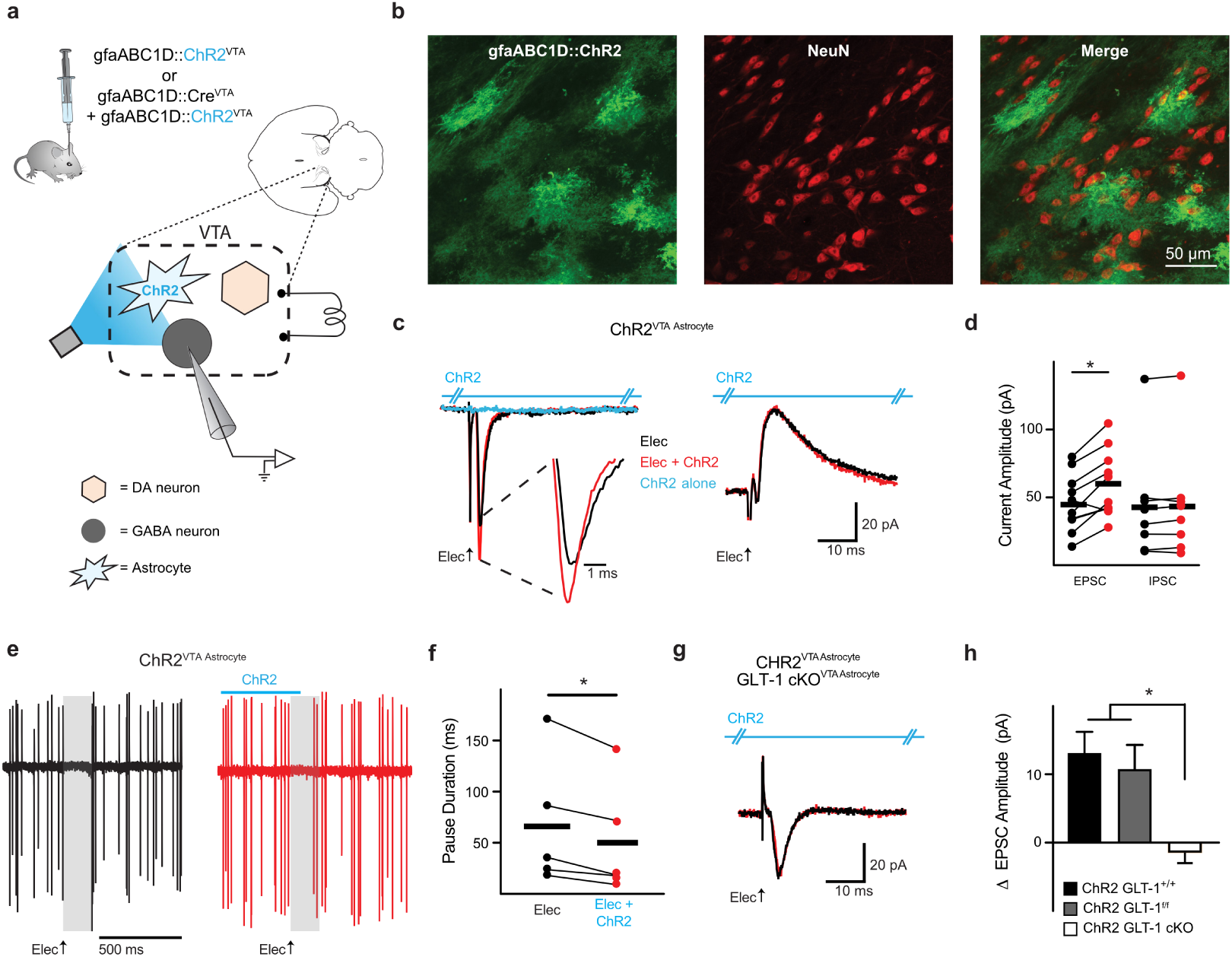
Astrocytic GLT-1 increases excitation of VTA GABA neurons. **(a)** Schematic showing the microcircuitry of the VTA and manipulations performed. (**b)** Immunocytochemistry images (20X objective) showing lack of co-localization between ChR2 (green, note characteristic astrocyte morphology) and NeuN (red) within VTA. (**c**) Example voltage-clamp recordings (left V_h_ = −55 mV, right V_h_ = +10 mV). The inward current (left), but not the outward current (right), increases when astrocytes are photoactivated (ChR2, blue hash marked horizontal line above traces) concurrently with electrical stimulation (Elec.) of neuronal afferents (red traces) in a GLT-1^+/+^ mouse. ChR2 photoactivation alone produces no response (blue trace; see Fig. S8). **(d)** Summarized data from VTA GABA neurons where ChR2 stimulation on VTA astrocytes increases peak EPSC amplitude (GLT-1^+/+^, paired t-test t_(9)_ = 4.457, P = 0.016, n = 10 cells) without altering IPSCs (paired t-test t_(7)_ = 1.318, P = 0.22, n = 8 cells, 4 mice). **(e)** 5 overlaid traces of a cell-attached recording from a VTA GABA neuron. Electrical stimulation of neuronal afferents in the VTA (black traces) elicits a pause in firing activity (grey area). However, when ChR2 on VTA astrocytes (gfaABC1D::ChR2^VTA^) is photoactivated concurrently with electrical stimulation (red traces) the pause duration is decreased (see spikes within grey area, which is pause duration of electrical stimulation alone). (**f)** Summarized data from GABA neurons demonstrating a decrease in pause duration (paired t-test t_(4)_ = 3.345, P = 0.028, n = 5 cells, 2 mice) elicited by ChR2 stimulation on VTA astrocytes. **(g)** The increase in EPSC amplitude is absent in a GLT-1 cKO^VTA Astrocyte^ mouse (GLT-1^f/f^, gfaABC1D::Cre^VTA^ + gfaABC1D::ChR2^VTA^). (**h**) Summarized data comparing the change in the EPSC amplitude between GLT-1^+/+^ (GLT-1^+/+^, gfaABC1D::ChR2^VTA^), GLT-1^f/f^ (GLT-1^f/f^, gfaABC1D::ChR2^VTA^), and GLT-1 cKO^VTA^ ^Astrocyte^ mice (One-way ANOVA, F_(2,22)_ = 7.352, P = 0.0036; post-hoc Tukey’s test GLT-1^+/+^ vs GLT-1^f/f^ P > 0.05, GLT-1^+/+^ vs GLT-1 cKO^VTA^ ^Astrocyte^ p < 0.05, GLT-1^f/f^ vs GLT-1 cKO^VTA^ ^Astrocyte^ P < 0.05, n = 11, n = 6, n = 8 cells respectively). **: P < 0.01, *: P < 0.05; Error bars indicate ± SEM.

We investigated whether local GABA neurons are subject to modulation by VTA astrocytes. We found that during voltage-clamp recordings of VTA GABA neurons (Fig. S4a-e), electrically-stimulated excitatory postsynaptic currents (EPSCs) increased in peak amplitude when the electrical stimulation was paired with ChR2 photoactivation on VTA astrocytes (GLT-1^f/f^, Elec = 22.0 ± 3.7 pA, Elec + ChR2 = 32.4 ± 6.6 pA; paired t-test t_(5)_ = 3.147, P = 0.025, n = 6 cells, 3 mice) with no effect on inhibitory postsynaptic currents (IPSCs) (GLT-1^f/f^, Elec = 44.8 ± 18.1 pA, Elec + ChR2 = 40.4 ± 22.1 pA; paired t-test t_(4)_ = 0.8756, P = 0.43, n = 5 cells, 3 mice; data not shown for GLT-1^f/f^ mice). When the same cells were recorded in cell-attached mode, electrical stimulation of all afferents in the brain slice (no blockers were present) elicited a net pause in firing. When the electrical stimulation was paired with photoactivation of astrocytes, the pause duration decreased, which is consistent with an increase in EPSC amplitude observed in voltage-clamp mode (Fig. 2e-f).

Because astrocyte photoactivation with ChR2 could influence glutamate transport ^28^,^30^,^35^,^36^, we next sought to determine whether VTA astrocytes increase excitation of local GABA neurons in a glutamate transporter-dependent manner. We drove a conditional knock out (cKO) by injecting an adeno-associated virus (AAV) into the VTA of GLT-1 floxed mice (GLT-1^f/f^) ^39^ to restrict recombination and loss of GLT-1 to VTA astrocytes (GLT-1^f/f^, AAV(gfaABC1D-Cre) → VTA; Fig. 2a, S6,7). The increase in EPSC amplitude elicited by astrocyte photoactivation was absent when GLT-1 was conditionally knocked out from VTA astrocytes (GLT-1 cKO^VTA^ ^Astrocyte^; Elec = 28.18 ± 4.5 pA, Elec + ChR2 = 28.6 ± 3.8; paired t-test t_(7)_ = 0.3152, P = 0.76, n = 8 cells, 3 mice; Fig. 2g), indicating that astrocyte activation increases EPSC amplitude in VTA GABA neurons via a mechanism that requires GLT-1 expression in astrocytes.

Contrary to what we observed from GABA neurons, when we recorded from VTA dopamine neurons (Fig. S4f-j), the IPSC peak amplitude increased, but the EPSC was unaffected, when electrical stimulation was paired with VTA astrocyte photoactivation (GLT-1^f/f^ IPSC, Elec = 44.7 ± 10.6 pA, Elec + ChR2 = 56.9 ± 11.4; paired t-test t_(6)_ = 4.243, P = 0.0054, n = 7 cells, 2 mice, GLT-1^f/f^ EPSC: Elec = 40.4 ± 11.3 pA, Elec + ChR2 = 42.8 ± 12.4; paired t-test t_(6)_ = 1.199, P = 0.27, n = 7 cells, 2 mice; data not shown for GLT-1^f/f^ mice). When the same dopamine cells were recorded in cell-attached mode, electrical stimulation of all afferents in the brain slice (no blockers present) also elicited a net pause in firing. However, when the electrical stimulation was paired with photoactivation of astrocytes, the pause duration increased (Fig. 3d-e), which is consistent with the increase in IPSC amplitude observed in voltage-clamp mode. The increase in IPSC amplitude was blocked when the glutamate transporter blocker, DHK (300 μM), was added during the same recording (paired t-test t_(6)_ = 5.675, P = 0.0013, n = 7 cells, 3 mice; Fig. 3f-g). Also, when GLT-1 was conditionally knocked out from VTA astrocytes, the IPSC amplitude on DA neurons was unaffected by astrocyte activation paired with electrical stimulation (GLT-1 cKO^VTA Astrocyte^; Elec = 51.7 ± 9.5 pA, Elec + ChR2 = 52.4 ± 10.2; paired t-test t_(7)_ = 0.2926, P = 0.77, n = 8 cells, 2 mice; Fig. 3h). Importantly, the effect of GLT-1 manipulation was dependent on active synaptic input since photoactivation of VTA astrocytes without electrical stimulation of neuronal afferents had no effect on either VTA dopamine or GABA neurons (Fig. 2c, 3b, S8). Thus, astrocytes use a GLT-1-dependent mechanism to increase both excitation in GABA neurons and inhibition in dopamine neurons, but only during concurrent activity of neuronal afferents.

**Figure 3.**
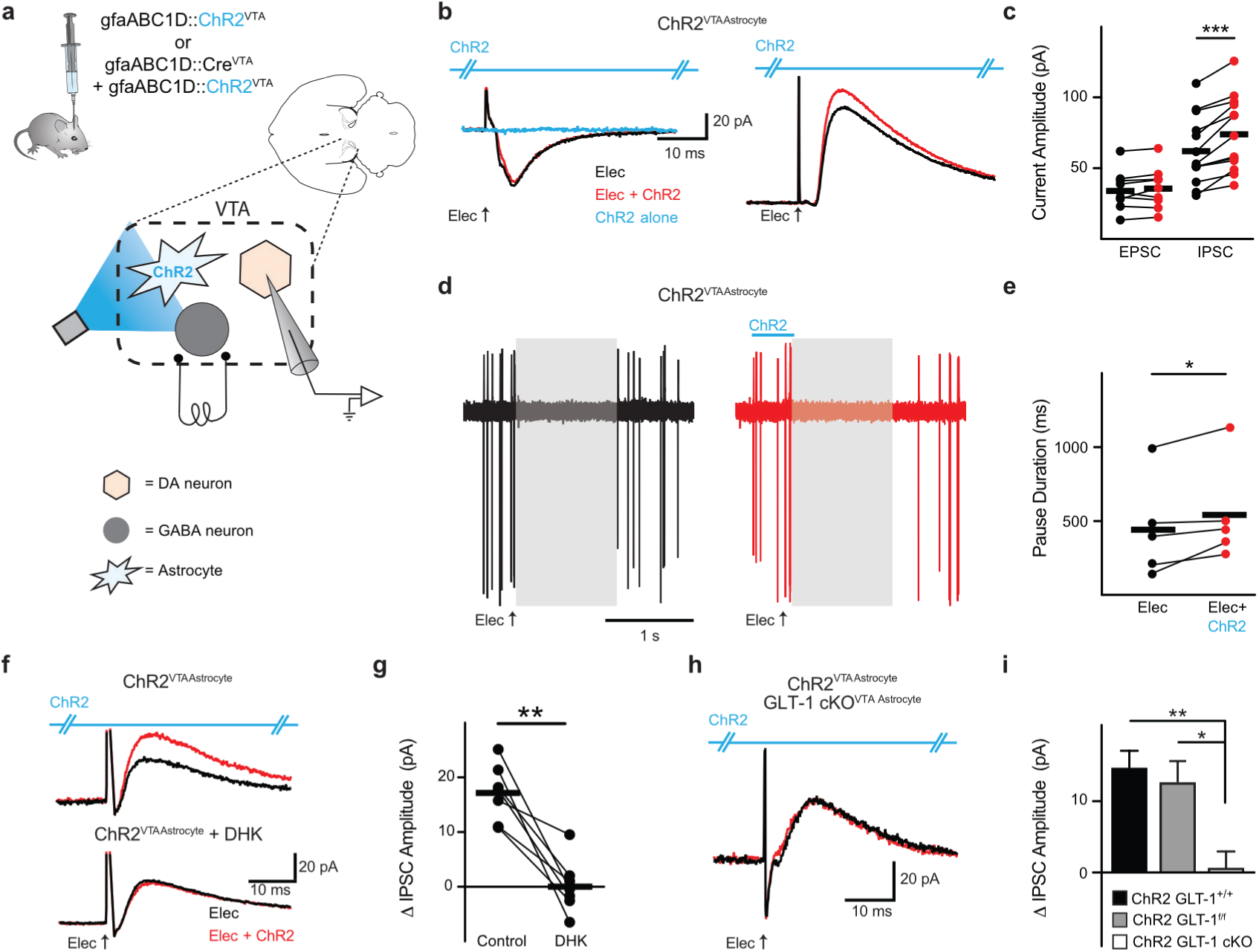
Astrocytic GLT-1 increases inhibition of VTA dopamine neurons. **(a)** Schematic showing the microcircuitry of the VTA and manipulations performed. **(b)** Example voltage-clamp recordings (left V_h_ = −55 mV, right (V_h_ = +10 mV). For dopamine neurons, the outward current (right), not the inward current (left), increases when ChR2 is photoactivated concurrently with electrical stimulation (Elec.) of neuronal afferents (red traces) in a GLT-1^+/+^ mouse. Photoactivation of ChR2 alone elicits no response (blue trace, and see Fig. S8). **(c)** Summarized data from VTA dopamine neurons where ChR2 stimulation on VTA astrocytes increases peak IPSC amplitude (GLT-1^+/+^, paired t-test t_(12)_ = 5.894, P = 0.0001, n = 13 cells, 8 mice) without altering EPSCs (paired t-test t_(9)_ = 1.129, P = 0.28, n = 10 cells, 8 mice). **(d)** 9 overlaid traces of a cell-attached recording and **(e)** summarized data from VTA dopamine neurons demonstrating an increase in pause duration (paired t-test t_(4)_ = 2.815, P = 0.04, n = 5 cells, 2 mice) when ChR2 stimulation on VTA astrocytes is delivered concurrently with electrical stimulation of neuronal afferents (red traces; grey areas are the pause duration following electrical stimulation alone). **(f, g)** The increase in IPSC amplitude (red trace, top) is abolished by pharmacologically blocking GLT-1 with 300 DHK (bottom traces). **(h)** When GLT-1 is conditionally knocked out from VTA astrocytes there is no change in IPSC caused by activating ChR2 on astrocytes. **(i)** Summarized data comparing the change in IPSC amplitude between GLT-1^+/+^ (GLT-1^+/+^, gfaABC1D:: ChR2^VTA^), GLT-1^f/f^ (GLT-1^f/f^, gfaABC1D:: ChR2^VTA^), GLT-1^f/f^, and GLT-1 cKO^VTA^ ^Astrocyte^ mice (One-way ANOVA, F_(2,27)_ = 7.732, P = 0.0022; post-hoc Tukey’s test GLT-1^+/+^ vs GLT-1^f/f^ P > 0.05, GLT-1^+/+^ vs GLT-1 cKO^VTA Astrocyte^ P < 0.05, GLT-1^f/f^ vs GLT-1 cKO^VTA Astrocyte^ P < 0.05, n = 11, n = 6, n = 8 cells respectively). **: P < 0.01, *: P < 0.05; Error bars indicate ± SEM.

### VTA GABA neurons mediate the astrocyte-dependent increase in inhibition of dopamine neurons

Given the surprising result of a glutamate transporter mediating the increase of an outward current in dopamine neurons, we next determined whether the outward current was indeed an IPSC mediated by an inhibitory input. We tested the sensitivity of the outward current to the GABAA receptor antagonist, bicuculline (30 μM). Both the electrically-induced IPSC, and the peak amplitude increase elicited by pairing electrical stimulation with photoactivation of astrocytes, were abolished by addition of bicuculline, indicating that all outward currents were mediated by GABAa receptor activation (paired t-test t_(4)_ = 3.682, P = 0.0015, n = 5 cells, 3 mice; Fig. 4a-c). However, only the increase in IPSC amplitude, not the electrically-elicited outward current per se, was sensitive to glutamate receptor antagonism (10 NBQX + 100 AP5, paired t-test t_(7)_ = 3.101, P = 0.0173, n = 8 cells, 4 mice, 4d-e). Since *1*) ChR2 activation increased the EPSC amplitude on VTA GABA neurons, and *2*) the increased IPSC on dopamine neurons was dependent on glutamate receptor activation, dopamine neurons may be inhibited via a circuit mechanism involving depolarization of GABA neurons.

**Figure 4.**
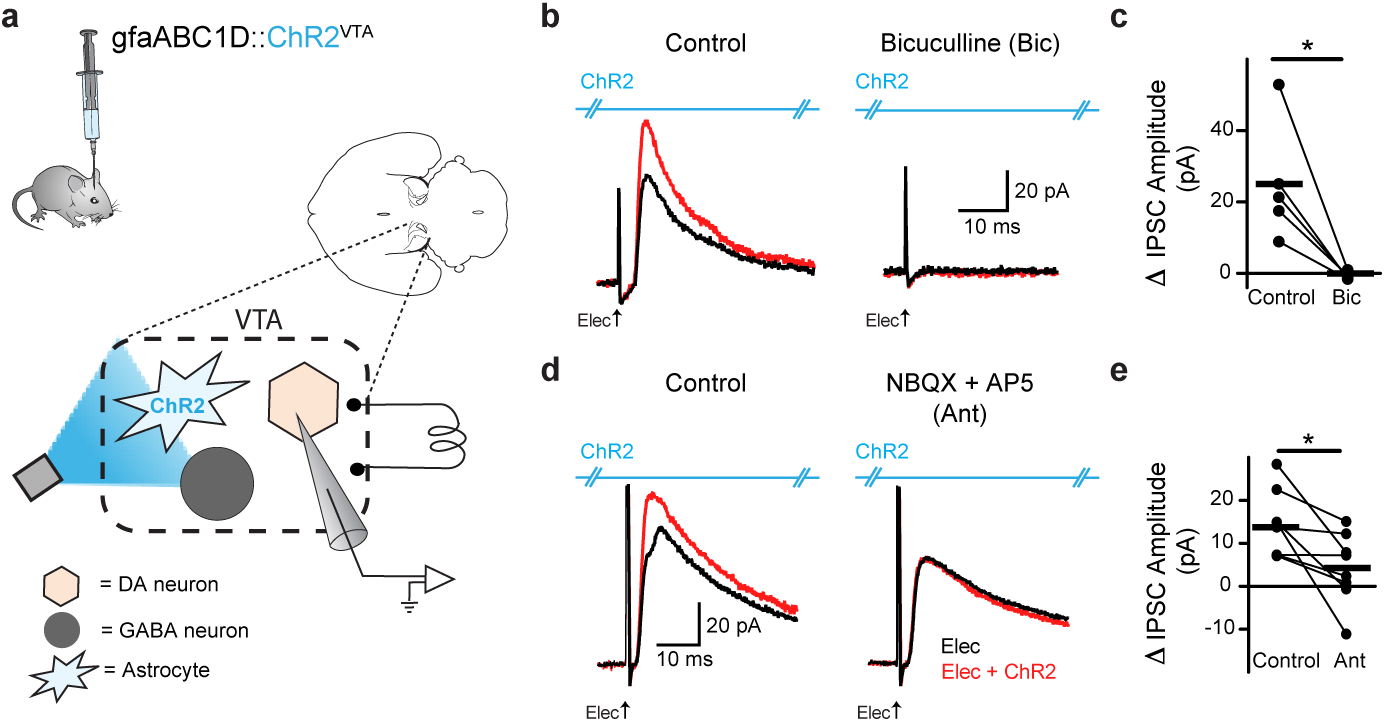
Increase in dopamine neuron inhibition is dependent on glutamate receptor activation. **(a)** Schematic showing the microcircuitry of the VTA and manipulations performed. **(b)** On the left is a voltage-clamp (Vh = +10 mV) recording from a VTA dopamine neuron where pairing electrical simulation (Elec., black traces) with ChR2 stimulation on VTA astrocytes (red traces) causes an increase in IPSC amplitude. On the right, when the GABAa antagonist, bicuculline (30 μM), is added during the same recording the IPSC is abolished. **(c)** Data summarizing the effect of bicuculline. **(d)** The increase in the IPSC amplitude was eliminated by glutamate receptor antagonists (10 μM NBQX + 100 μM AP-5) with no effect on the amplitude of the IPSC elicited by electrical stimulation alone (black traces) (P = 0.99, n = 8 cells, 4 mice). **(e)** Data summarizing the effect of glutamate receptor antagonists. *: P < 0.05.

To directly test whether local GABA neurons mediated the effects of astrocyte activation on dopamine neuron inhibition, we optogenetically hyperpolarized VTA GABA neurons (VGAT^Cre^::eNpHR^VTA^) during photoactivation of ChR2 on VTA astrocytes (gfaABC1D::ChR2^VTA^; Fig. 5a-b; S9). Optogenetic hyperpolarization of VTA GABA neurons prevented the increase in IPSC amplitude, recorded from neighboring dopamine neurons, that resulted from combining electrical stimulation with optogenetic activation of VTA astrocytes during the same recordings (unpaired t-test t_(18)_ = 3.961, P = 0.0009, electrical + ChR2: n = 7 cells, 4 mice, electrical + ChR2 + NpHR: n = 13 cells, 6 mice; Fig. 5c-d).

**Fig. 5.**
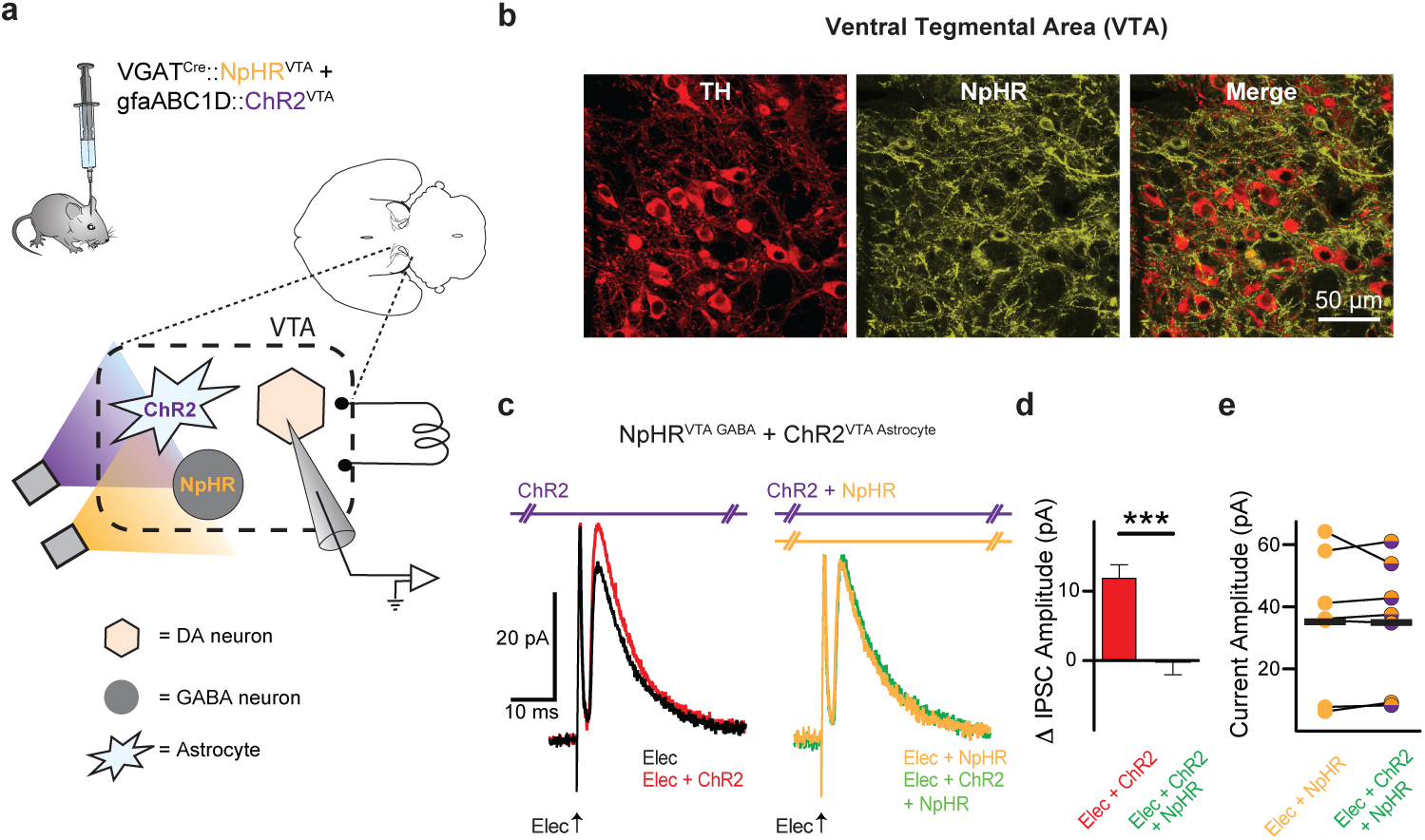
Astrocyte-dependent increase in dopamine neuron inhibition is mediated by VTA GABA neurons. **(a)** Experimental manipulation. **(b)** Immunocytochemistry demonstrating TH positive cells (red) and eNpHR expression (yellow). On the right are the two images merged together. (**c)** Voltage-clamp traces (V_h_ = +10 mV) from a VTA dopamine neuron. ChR2 stimulation during electrical stimulation (Elec + ChR2, red trace; ChR2 stimulated at 405 nm to minimize activation of eNpHR) increases the IPSC amplitude relative to electrical stimulation alone (Elec, black trace). When ChR2 on astrocytes is activated concurrently with eNpHR activation on GABA neurons during electrical stimulation (Elec + ChR2 + eNpHR, green trace) the increase in IPSC amplitude was eliminated. Electrical stimulation with GABA neuron hyperpolarization alone (Elec + eNpHR, yellow trace) has no difference relative to electrical stimulation along with ChR2 on astrocytes and eNpHR activation on GABA neurons (Elec + ChR2 + eNpHR, green trace). (**d)** The increase in IPSC amplitude when ChR2 was photoactivated with electrical stimulation (Elec + ChR2, red) was blocked when GABA neurons were simultaneously hyperpolarized (Elec + eNpHR + ChR2, green). **(e)** IPSC amplitudes during electrical stimulation with GABA neuron hyperpolarization (Elec + eNpHR, yellow) were not changed when ChR2 stimulation was added (Elec + eNpHR + ChR2, purple-yellow; paired t-test t_(6)_ = 0.1247,P = 0.90, n = 7 cells, 4 mice). ***: P < 0.001; Error bars indicate ± SEM.

### VTA astrocytes drive real-time avoidance mediated by local GABA neurons

Previous studies demonstrate that direct optogenetic depolarization of VTA GABA neurons inhibits the activity of neighboring dopamine neurons and elicits avoidance behavior ^6^,^7^. Our recordings demonstrate that VTA astrocytes can increase the response of GABA neurons to active excitatory afferent input, leading to increased inhibition of dopamine neurons. We therefore aimed to determine whether ChR2 activation of astrocytes would directly drive avoidance behavior. Extending our same optogenetic strategy to behavioral assays, we found that photoactivation of VTA astrocytes induced real-time avoidance in control mice in a real-time place preference (RTPP) assay (GLT-1^+/+^, paired t-test t_(17)_ = 11.56, P = 0.0001, n = 18 mice; Fig. 6b, e; S10; Supplementary Video 1). The effect of astrocyte activation was reversible since mice could be manipulated to avoid either side of the chamber by changing which side was paired with laser stimulation (Fig. 6b-c). The avoidance behavior was also dependent on astrocytic GLT-1 since conditional knockout restricted to VTA astrocytes (GLT-1 cKO^VTA Astrocyte^, gfaABC1D::ChR2^VTA^) eliminated the real-time avoidance elicited by photoactivating VTA astrocytes (paired t-test t_(10)_ = 0.1399, P = 0.98, n = 11 mice; Fig. 6d-e).

**Figure 6.**
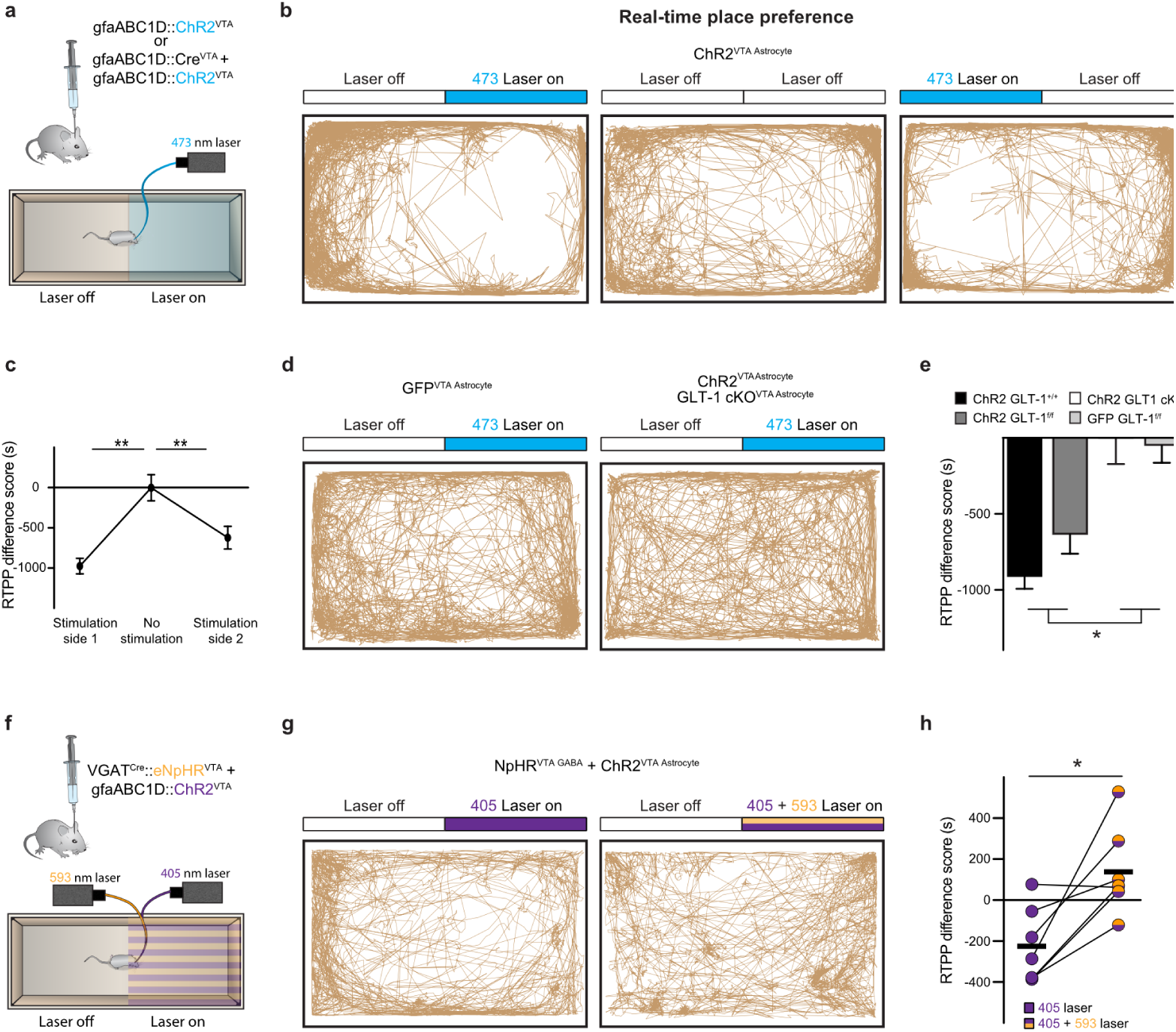
Real-time avoidance is dependent on VTA astrocytes and mediated by local GABA neurons. **(a)** Experimental manipulations for panels b-e. (**b)** Three track plots from one ChR2^VTA Astrocyte^ mouse (GLT-1^+/+^, gfaABC1D::ChR2^VTA^). (left) The mouse avoided the side of the chamber where ChR2 was stimulated in the VTA (473 nm laser on). When the laser was left off both sides (Laser off) the mouse showed no avoidance. (right) With laser stimulation on the opposite side, the mouse then avoided the opposite side. **(c)** Summarized data showing avoidance of the laser-paired side (Repeated Measures ANOVA, F_(1.413,11.3)_ = 18.43, P = 0.0006; post-hoc Tukey’s test, stimulation side 1 vs no stimulation P < 0.05, stimulation side 1 vs stimulation side 2 P > 0.05, Stimulation side 2 vs no stimulation P < 0.05, n = 9 mice). **(d)** (left) Track plot from a mouse expressing GFP on VTA astrocytes (GFP^VTA^ ^Astrocyte^; GLT-1^f/f^, gfaABC1D::GFP^VTA^) shows no avoidance. (Right) track plot from a mouse expressing ChR2 on, but with GLT-1 conditionally knocked out from, VTA astrocytes (GLT-1 cKO^VTA Astrocyte^; GLT-1^f/f^, gfaABC1D::ChR2^VTA^ + gfaABC1D::Cre^VTA^) showing no avoidance. **(e)** Summarized data showing avoidance elicited by ChR2 activation in GLT-1^+/+^ and GLT-1^f/f^ mice. However, loss of GLT-1 from VTA astrocytes eliminates avoidance (One-way ANOVA, F_(3,49)_ = 16.11, P = 0.0001; post-hoc Tukey’s test, GLT-1^f/f^ relative to: GLT-1 cKO^VTA^ ^Astrocyte^ p < 0.05, GLT-1^+/+^ P > 0.05, GFP P < 0.05; cKO^VTA^ ^Astrocyte^ relative to: GLT-1^+/+^ P > 0.05, GFP P < 0.05; GLT-1^+/+^ P < 0.05). (**f**) Experimental manipulations for panels g-h. **(g)** Track plots from a VGAT^CRE^::eNpHR^VTA^ mouse with ChR2 stimulation on VTA astrocytes (eNpHR^VTA GABA^ + ChR2^VTA Astrocyte^). Avoidance observed during ChR2 stimulation (left; ChR2 stimulated at 405 nm to minimize activation of eNpHR) was blocked when astrocytes were photoactivated concurrently with GABA neuron hyperpolarization (right). **(h)** Summarized data demonstrating GABA neuron hyperpolarization blocks avoidance elicited by ChR2 on VTA astrocytes. ***: P < 0.001 **: P < 0.01, *: P < 0.05; Error bars indicate ± SEM.

We then confirmed that GABA neuron depolarization is necessary for eliciting real-time place avoidance by optogenetically hyperpolarizing VTA GABA neurons concurrently with ChR2 activation of VTA astrocytes. Similar to the increase in IPSC amplitude in dopamine neurons, real-time avoidance elicited by photoactivation of VTA astrocytes from VGAT^Cre^::NpHR^VTA^ mice was blocked by concurrently hyperpolarizing local GABA neurons (paired t-test t_(6)_ = 3.642, eNpHR + ChR2; P = 0.01, n = 7 mice; Fig. 6g-h, S9). Therefore, astrocyte activation reliably induced real-time avoidance behavior through astrocytic GLT-1-dependent and GABA neuron depolarization-dependent mechanisms that are consistent with the circuit pathway established in our ex vivo experiments.

### Learned avoidance, but not approach, is dependent on VTA astrocytes

Because of the VTA’s well known involvement in cue-outcome associative learning, we next sought to determine if astrocyte manipulation-evoked avoidance was sufficient to serve as a motivationally relevant signal in an associative learning context. The place conditioning paradigm is particularly well-suited for examining approach or avoidance since it makes use of the fact that mice will readily learn to approach or avoid distinct environments previously paired with either rewarding or aversive events, respectively ^40^. We implemented a conditioned place avoidance (CPA) paradigm using photoactivation of VTA astrocytes as a conditioning stimulus paired to a contextual cue (chamber) during conditioning sessions. Astrocyte activation was sufficient to elicit learned avoidance in the CPA assay (gfaABC1D::ChR2^VTA^; GLT-1^f/f^: paired t-test t_(9)_ = 3.604, P = 0.0057, n = 10 mice; Fig. 7b, d; S10), while GLT-1 cKO restricted to VTA astrocytes eliminated expression of avoidance learning during the assay (GLT-1 cKO^VTA Astrocyte^, gfaABC1D::ChR2^VTA^: paired t-test t_(11)_ = 0.7296, P = 0.48, n = 12; Fig. 7c-d). Thus, VTA astrocytes mediate associative learning of aversive cues through a mechanism that requires expression of astrocyte GLT-1.

**Figure 7.**
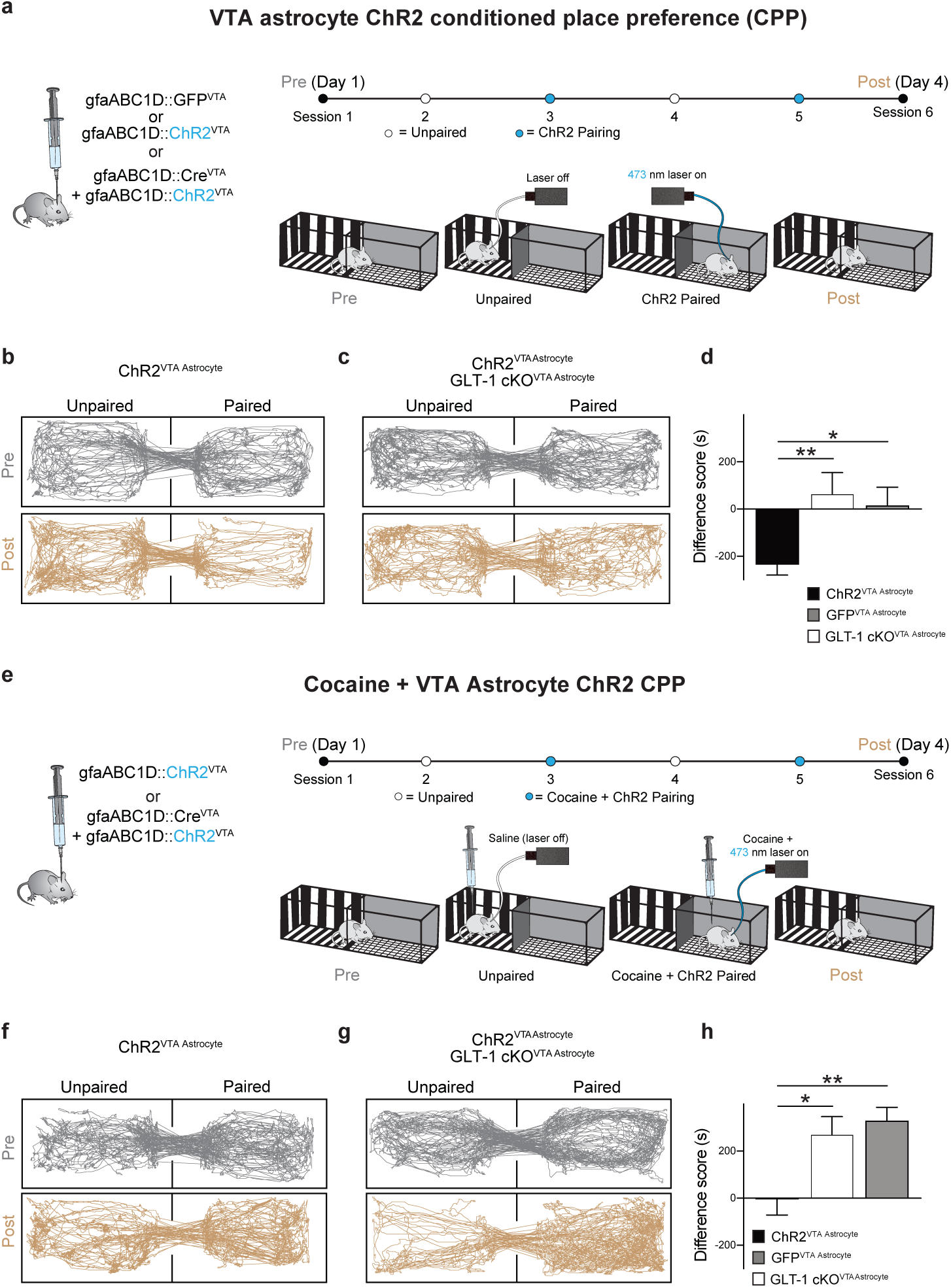
VTA Astrocytes mediate learned avoidance and prevent learned preference for reward. **(a)** Experimental design and manipulations for panels b-d. **(b)** Track plots showing that conditioning with ChR2 photoactivation on VTA astrocytes elicits learned avoidance in a GLT-1^+/+^ mouse (gfaABC1D::ChR2^VTA^). **(c)** The learned avoidance is absent in a mouse where GLT-1 was removed from VTA astrocytes (gfaABC1D:: ChR2^VTA^, GLT-1 cKO^VTA Astrocyte^). **(d)** Comparing the difference scores in mice expressing either ChR2 (GLT-1^+/+^ and GLT-1^f/f^, GLT-1 cKOVTA Astrocyte) or GFP (gfaABC1D:: ChR2^GFP^) in VTA astrocytes. **(e)** Experimental design and manipulations for panels f-h. **(f)** Track plots showing astrocyte activation paired with cocaine administration does not elicit either learned avoidance or preference in a GLT-1^f/f^ mouse (gfaABC1D::ChR2^VTA^). **(g)** However, when GLT-1 is removed from VTA astrocytes (gfaABC1D::ChR2^VTA^, GLT-1 cKO^VTA Astrocyte^) a mouse displays robust learned preference for VTA astrocyte ChR2 photoactivation paired with cocaine administration. **(h)** Comparing the difference scores in mice expressing either ChR2 (GLT-1^+/+^ and GLT-1^f/f^, GLT-1 cKO^VTA Astrocyte^) or GFP (gfaABC1D::ChR2^GFP^) in VTA astrocytes. **: P < 0.01, *: P < 0.05; Error bars indicate ± SEM.

When a cue predicts both reward and aversion, dopamine neurons’ activity responses reflect the integrated value of both outcomes ^9^, indicating a convergence of information at or before the level of the dopamine neuron. As a step in establishing astrocytes as functional elements within the VTA approach-avoidance circuit, we sought to disambiguate whether GLT-1 on astrocytes is positioned within a functional pathway specific to avoidance, or a common pathway that mediates expression of both avoidance and preference. If astrocytes mediate a common pathway, absence of astrocyte GLT-1 would be expected to abolish expression of both approach and avoidance. To dissect pathway specificity or generality, we used a conditioned place preference (CPP) paradigm to pair a contextual cue with a conditioning stimulus that contained both a strong reward (cocaine, 15 mg/kg) and aversion (photoactivation of VTA astrocytes). Consistent with prior work ^41^, cocaine alone drove robust approach in the CPP assay (GLT-1^+/+^ GFP: paired t-test t_(8)_ = 3.735, n = 9; P = 0.0057; not shown). However, when cocaine was paired with ChR2 activation on VTA astrocytes during the conditioning sessions, both the preference for cocaine and the avoidance of ChR2 photoactivation were abolished (gfaABC1D::ChR2^VTA^: paired t-test t_(7)_ = 0.7588, P = 0.47, n = 8 mice, Fig. 7f, h, S10). Since expression of CPA was mediated by GLT-1 on VTA astrocytes we next sought to determine whether cocaine CPP was also dependent on VTA astrocyte GLT-1 expression. However, preference for cocaine remained normal even when cocaine was paired with photoactivation of VTA astrocytes during conditioning in GLT-1 cKO^VTA Astrocyte^ mice (One-way ANOVA, F_(2,33)_ = 8.295, P = 0.0012; post-hoc Tukey’s test GFP vs ChR2 P > 0.05, GFP vs GLT-1 cKO^VTA Astrocyte^ P < 0.05, ChR2 vs cKO^VTA Astrocyte^ P > 0.05; ChR2: n = 14, GFP: n = 10, GLT-1 cKO^VTA^ ^Astrocyte^ n = 7, Fig. 7g-h, S10). Thus, although CPA requires GLT-1 expression in VTA astrocytes and can suppress expression of learned approach, CPP for cocaine is independent of astrocytic GLT-1 in the VTA.

### Astrocytic GLT-1 in the VTA is necessary for endogenous approach-avoidance behavior

To determine the impact of VTA astrocytic GLT-1 function on endogenous behavior we tested whether selective deletion of GLT-1 in VTA astrocytes is sufficient to skew an animal’s natural behavior in an open field test as an ethological assay that indexes approach-avoidance conflict. Mice with GLT-1 cKO restricted to VTA astrocytes (GLT-1 cKO^VTA Astrocyte^) displayed a decreased tendency to remain in the peripheral zone near the walls of the chamber, and increased tendency to explore the center of the open field, relative to littermate controls (unpaired t-test t_(16)_ = 2.449, P = 0.026, n = 9 mice, Fig. 8b-c). Disrupted approach-avoidance behavior in GLT-1 cKO^VTA^ ^Astrocyte^ mice indicates that VTA astrocytes are an integral component of a circuit within the VTA capable of coordinating approach and avoidance behaviors.

**Figure 8.**
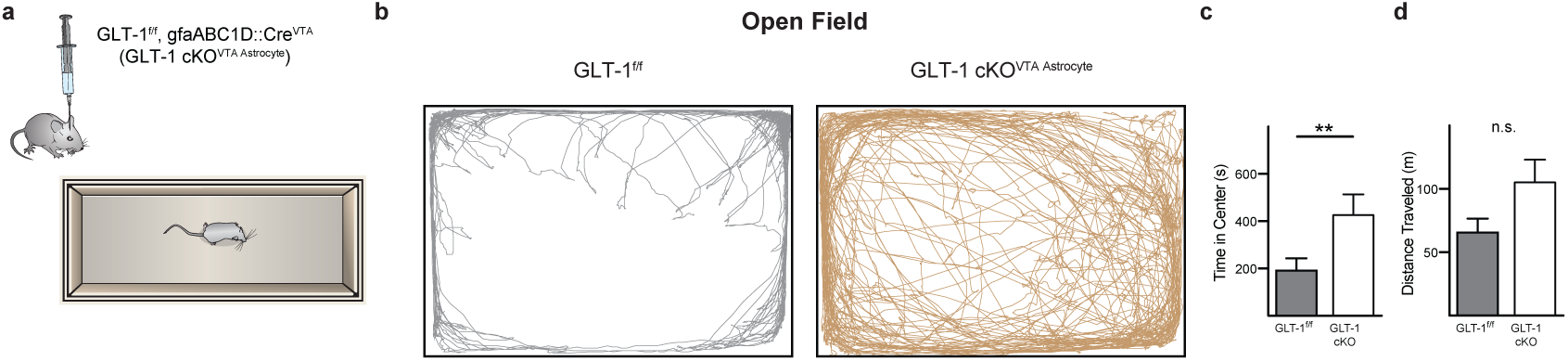
GLT-1 removal from VTA astrocytes disrupts endogenous avoidance behavior. **(a)** Schematic depicting experimental design. **(b)** Track plots from a GLT-1^f/f^ mouse with only a fluorescent reporter (GFP) on astrocytes displaying normal thigmotaxis in an open field (left), and a GLT-1 cKO^VTA Astrocyte^ (GLT-1^f/f^, gfaABC1D::Cre^VTA^) mouse with increased tendency to explore the center of the open field (right). (**c, d)** GLT-1 cKO^VTA Astrocyte^ mice spent more time exploring the center of the open field without affecting locomotor activity (**d**; unpaired t-test t_(16)_ = 1.981, P = 0.065, n = 9 mice). **: p < 0.01; Error bars indicate ± SEM.

## DISCUSSION

### Astrocytes control signaling for avoidance

Our experiments identify, through a series of genetic and optogenetic strategies in slice and in vivo, a specialized VTA microcircuit that is regulated by astrocytes to drive avoidance behavior. We observed that local astrocytes within the VTA circuit modulate glutamate availability via GLT-1 to dynamically regulate active GABAergic inhibition of dopamine neurons (Fig. 9). As a result, VTA astrocytes induced robust, time-locked behavioral avoidance de novo. Rapid manipulation of astrocyte glutamate transport in the VTA served as a powerful stimulus that, when paired with cocaine, was sufficient to impede behavioral expression of cocaine CPP. Thus, despite cocaine’s many systemic effects, local astrocyte manipulation within the VTA was sufficient to override expression of preference for a highly salient cocaine-associated cue. These experiments suggest that midbrain astrocytes have the capacity to drive behavior, and not merely modulate the gain or polarity of an ongoing behavior. They underscore a computational role for astrocytes in the VTA network, and provide the first evidence that VTA astrocytes can selectively and reliably determine complex motivated behavior.

**Figure 9.**
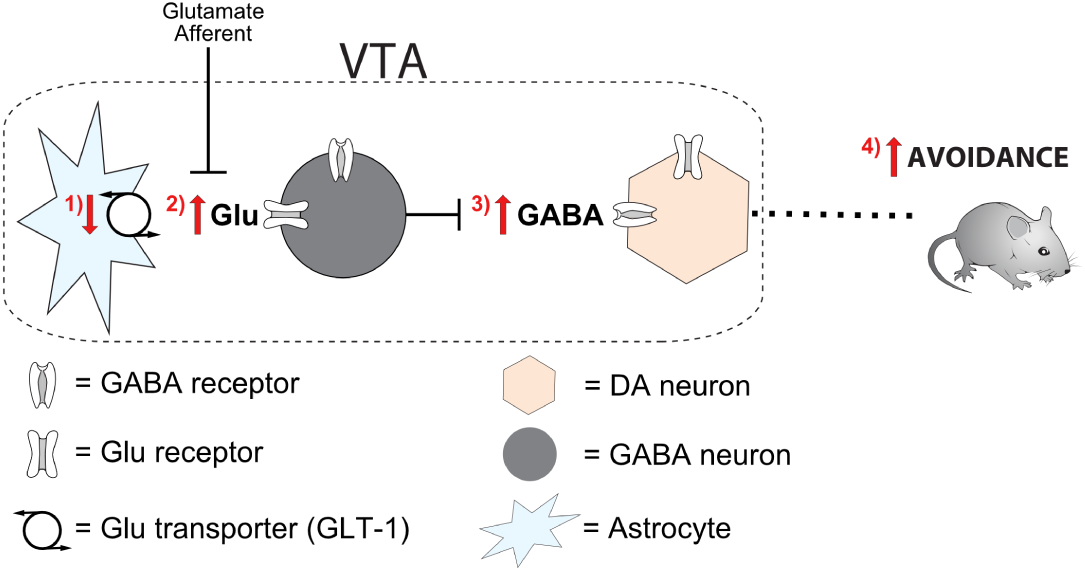
A neuron-astrocyte circuit for avoidance. In a sequence of events, (1) reduction of glutamate uptake by the astrocytic glutamate transporter, GLT-1, leads to (2) increased glutamate concentration at synapses on VTA GABA neurons. (3) The resultant increase in GABA release at synapses on DA neurons inhibits DA neuron activity, eliciting (4) real time and learned avoidance behavior.

### Separable circuits for avoidance and approach in the VTA

The complexity of afferents arriving at the VTA points to its importance as a nexus where signals related to sensory experience, valence, and behavioral state intersect with associative processes to cue motivated behavior. For example, separate projections from the bed nucleus of the stria terminalis (BNST) to the VTA mediate both aversive and rewarding signals that promote and attenuate anxiety phenotypes, respectively ^42^. But, the BNST provides equivalent projections to both GABA and dopamine neurons within the VTA ^43^. Afferents from the pedunculopontine nucleus to both VTA GABA and dopamine neurons mediate approach behaviors like conditioned place preference for reward ^44^,^45^. The VTA is therefore capable of parsing avoidance signals from different afferents to drive avoidance behavior and appropriately coordinate approach and avoidance. But a distinct circuit for avoidance capable of sufficiently opposing approach had not been described.

Inhibition exerts a powerful regulatory influence on dopamine neurons ^46^, and prior studies establish that direct depolarization of VTA GABA neurons elicits real-time avoidance but fails to disrupt conditioned responses to reward-predictive cues ^6,7^. Our experiments reliably reproduce the real-time avoidance results solely through optogenetic activation of VTA astrocytes. Furthermore, they illustrate that a single astrocytic mechanism can facilitate dopamine neuron inhibition, induces both active and learned avoidance, and interferes with expression of cocaine CPP. All rely on GLT-1 expression at VTA astrocytes; when VTA astrocytes lack functional GLT-1, laser-cued conditioning fails to produce learned avoidance, and also fails to block expression of cocaine CPP (Fig. 7g-h). The selective influence of astrocyte mechanisms on avoidance but not approach in the dopaminergic VTA circuit reinforces the view that the microcircuits mediating avoidance and approach are functionally separate within the VTA, but converge at the level of inputs to dopamine neurons ^9^.

Our results demonstrate that VTA astrocytes can exert opposing effects on GABA and dopamine neurons through serial excitation of GABA neurons and subsequent inhibition of dopamine neurons (Fig. 9). The unexpected lack of direct astrocyte-mediated glutamatergic effects at dopamine neurons concurrent with those observed at GABA neurons highlight the separate circuits for avoidance and approach, and may reflect a partitioning mechanism akin to the basal ganglia wherein distinct subpopulations of astrocytes are functionally linked to either D1-type or D2-type receptor-expressing populations of spiny projection neurons ^17^ (but see also ^47^). One possibility is that astrocytic glutamate transport is differentially recruited by GABA and dopamine neurons in the VTA. Astrocyte populations that express distinct complements of transporters and receptors have been reported in the hippocampus ^16^,^48^, and may similarly tile the VTA to partition neuron types and their synapses. Additionally, GLT-1 is expressed in both astrocytes and neurons ^49^. Conditional genetic deletion of GLT-1 from neurons blunts the effects of amphetamine, and, in direct contrast to the VTA astrocyte GLT-1 knockout (Fig. 8), has no effect on open field behavior ^50^. Characterizing the molecular diversity, morphology and distribution of astrocytes within the VTA will be required to fully mine the dimensions of their computational capacity ^51^.

### Optogenetic manipulation of astrocytes

Astrocytes discriminate the activity of discrete synapses originating from different afferents ^14^,^15^,^20^. Recent work points to the unique molecular and physiological profiles of ventral midbrain astrocytes relative to their telencephalic counterparts ^51^, highlighting the need for approaches tailored to suit the functional dynamics of the system under study ^52^. With this in mind, we implemented an astrocyte manipulation protocol aligned to the temporal dynamics of VTA dopamine neurons (where bursting occurs on the order of hundreds of milliseconds) and associated behaviors. Expression of ChR2 specifically in VTA astrocytes exploits the tight association between astrocyte cation permeability and GLT-1 function ^25–30,35,37^. By engaging chiefly ion gradients, optical stimulation recruits native astrocyte machinery in electrochemically realistic processes. This approach succeeds in expressing astrocyte-specific, temporally controlled dynamics that make few assumptions about the relatively unknown cellular processes of VTA astrocytes ^34^. Finally, the use of visible light wavelengths minimizes reactive gliosis, which often confounds modulatory dynamics with long lasting inflammatory processes ^53^. It is notable that in vivo astrocyte manipulation produced real-time aversion that was readily reversed and spatially manipulated (Fig. 6b-c), signifying that ChR2 activation of astrocytes reflects dynamic modulation rather than long-lasting reactive processes. Optogenetic manipulation of astrocytes also affects only those afferents that are active during ChR2 photoactivation (Fig. 2–3, S8) ^14^. Therefore, during our conditioned place paradigms only those neuronal inputs that are active during the paired conditioning session will be facilitated during laser activation.

### Implications for pathophysiologies

As a core symptom of anxiety disorders, maladaptive avoidance is considered to be an underlying mechanism maintaining anxiety ^1^. Avoidance is a decision to refrain from potential negative outcomes at the expense of potential rewards. It is in this context that our finding is most relevant; VTA astrocyte GLT-1 can be manipulated to change inhibitory dynamics on dopamine neurons that interfere with the expression of conditioned place preference for reward. It is significant that the conditional loss of GLT-1 eradicates dynamical control of inhibition in the VTA and concurrently reduces avoidance behavior, but preserves approach to reward. Thus, separable pathways within the VTA drive behavioral responses to reward and aversion ^9^, and may be subject to distinct pathophysiologies that culminate in maladaptive learning processes ^1^. VTA astrocytes may therefore reflect a specific target for therapeutics, which have the added advantage of producing effects limited to activated synapses.

## CONCLUSIONS

Our results establish a previously unknown, intra-VTA neuron-astrocyte circuit mechanism specific to avoidance behavior through which astrocytes exert fast, dynamic control of dopamine neuron firing by modulating afferent glutamatergic drive on GABA neurons (glutamatergic afferents → astrocyte GLT-1 → GABA neuron EPSP→· dopamine neuron IPSP → avoidance; Fig. 9). We conclude that VTA astrocytes orchestrate temporal control of dopamine neuron firing, rendering them a decisive element in the circuit computation of avoidance, and promising targets for new therapeutic strategies to intervene in pathological disorders of behavior.

## METHODS

### Animals

Male and female C57BL/6J mice, Vgat-ires-Cre mice (016962 Stock) and GLT-1^f/f^ (GLT-1^flox/flox^) ^39^ mice were used. GLT-1^f/f^ mice (Slc1A2tm1.1Pros; MGI: 5752263) were obtained from the founder colony at Boston Children's Hospital. All procedures were approved by the University of Texas at San Antonio Institutional Animal care and Use Committees in accordance with the National Institutes of Health guidelines.

### Creation of AAV-TAS-gfaABC1D-ChR2-EYFP

With the creation of an abbreviated GFAP promoter that maintained astrocyte specificity ^38^, expression is amplified by using a transcriptional amplification strategy (TAS). This includes cloning a mini-CMV promoter driving expression of Gal4 binding domain fused with the p65 transcriptional activation domain on the opposite strand, and 5 concatenated Gal4 binding sites between the two promoters, mini-CMV and gfaABC1D ^54^. Using PCR, the TAS-gfaABC1D enhancer and promoter was amplified and isolated with AscI/SalI sites on the 5' and 3' ends, respectively. This fragment was subcloned into a AAV-CaMKII-ChR2-EYFP by digesting the plasmid with MluI/SalI to remove the CaMKII promoter and have compatible cohesive ends with the new promoter. The AAV vector was packaged and pseudo-typed with AAV5 at the UNC vector core.

### Stereotaxic injections and fiber optic implantations

8 to 12 week old mice were anesthetized using isoflurane anesthesia. Stereotaxic injections of different viruses (250 nL at a rate of 100 nl/min) were delivered to the VTA using the following coordinates relative to bregma AP: −2.6 to −3.2 ML: ±0.6 DV: −4.5. Mice received either AAV5/TAS-gfaABC1D-ChR2-eYFP or AAV5-GFA104-eGFP. Vgat-ires-Cre mice received injections of AAV5/gfaABC1D-ChR2-eYFP and AAV5/EF1α-DIO-eNpHR3.0-eYFP. GLT-1^f/f^ mice received injections of AAV5/gfaABC1D-ChR2-eYFP only or AAV5/gfaABC1D-ChR2-eYFP with AAV-gfaABC1D-Cre.

All mice were implanted with custom made fiber optic cannulas 100 μm above the injection site. Custom made cannulas were constructed using 1.25 mm diameter ferrule, 200 μm fiber optic cable with NA 0.39 and a two-part epoxy. The fiber optic cannulas were secured to the skull of each mouse using miniature screws and dental cement. Mice were singly housed and allowed to recover for 12 days before behavioral experiments were performed. A total of 3 mice were excluded from analysis for having virus and/or fiber optics outside of the VTA.

### Immunohistochemistry

Electrophysiology experiments: Once recordings were completed, the slices were incubated in 4% paraformaldehyde (PFA) for 30 minutes and moved to phosphate buffered solution (PBS) with .02% sodium azide for long-term storage at 4°C.

Mice used for behavioral experiments were anesthetized and intracardially perfused with ice-cold PBS, followed by 4% PFA. The brains were removed and placed in 4% PFA overnight at 4°C. The brains were next rinsed with PBS and 100 μM coronal slices containing VTA were collected. The slices were placed in PBS with .02% sodium azide for long-term storage at 4°C.

On day one slices were rinsed twice with PBS for ten minutes. Slices were blocked and permeabilized using 5% normal goat serum (NGS)/0.2% triton X-100 in PBS for two hours at room temperature. Slices were incubated overnight at 4°C with the following primary antibodies: Rabbit anti NeuN (1:1000, ABCAM), chicken anti tyrosine hydroxylase (1:500, AB9702), rabbit anti EAAT-2 (1:500, AB41621), mouse anti GABA (A0310) in 1% normal goat serum/0.1% triton X-100 with PBS. On day two, slices were washed three times, for ten minutes with PBS/0.05% triton X-100. Afterwards the slices were incubated with the following secondary antibodies: goat anti-rabbit Alexa 488 (1:1000, A11034) or Alexa 546 (1:1000, A11035), goat antichicken Alexa 488 (1:1000, A11039), Alexa 546 (1:1000, A11040) or 647 (1:1000, A21449) in 1% normal goat serum/0.1% triton X-100 in PBS for 2 hours at room temperature. Finally, the slices were washed three times for ten minutes with PBS/0.05% triton X-100 at room temperature and mounted onto slides using Fluoromount-G (0100–20). Bright field and fluorescence images for fiber and opsin placement were obtained using a CCD camera on a fluorescent stereoscope. Sections containing the ventral midbrain were imaged using a Zeiss confocal at 4 and 20X magnification with 1.0 μm thick optical sections. Each confocal image was obtained using identical settings for gain, laser and pinhole size. Images were then analyzed using ImageJ FIJI software.

### Flow Cytometry

#### Dissociation

The VTA was carefully sectioned from GLT-1^f/f^ and cKO mice in 1X Hanks’s Balanced Salt Solution (HBSS). The tissue was minced and incubated in pronase and repeatedly triturated every 20 minutes for 1 hour. DNASE (100 units/ml, Worthington) was added to the HBSS containing pronase solution to prevent cell aggregation. After 1 hour the cell suspension was diluted with 1X HBSS and centrifuged at 500g for 5 minutes. The pellet was resuspended in HBSS and filtered through a 100 μm filter. The cells were counted, and the filtrate was centrifuged and resuspended in Phosphate Buffer Saline (PBS).

#### Fixing and Staining

The cells were resuspended in 4% paraformaldehyde (PFA; Sigma) for 15 minutes and subsequently washed with 100 mM glycine (Sigma) in PBS to remove any traces of PFA. After centrifugation, the cells were resuspended in 5% donkey serum in 0.3% Triton-X for 30 minutes to block and permeabilize the cell membrane. The cells were then incubated with primary antibodies: mouse anti-GFAP (1:1000, G3893), rabbit anti EEAT-2 (1:1000, AB41621) followed by secondary antibodies: donkey anti mouse Alexa 488 (1:1000, A21202), donkey anti rabbit Alexa 555 (1:1000, A31572) for 20 and 15 minutes, respectively.

#### Analysis

Cells were analyzed in a BD LSR II flow equipped with three excitation lasers (488 nm, 555 nm and 633 nm). Briefly, events were selected based on their scatter properties indicative of cells, then selected for single cells, and selected based on fluorescence intensities. All gates were determined based on unstained, single-color and fluorescent minus 1 GLT-1^f/f^ of mouse brain cells. Gating analysis was done using FlowJo software (Treestar).

### Behavior

#### Open field test

Mice were placed in a custom-made plastic rectangular box (50 cm X 30 cm) and the box was divided into a center and outer zone with the outer zone being 7 cm from the wall. Mice were allowed to freely explore for 30 minutes. Anymaze (Stoelting) video tracking system was used to track the distance each mouse traveled and how much time they spent in each zone for all behavioral experiments.

#### Real-time place preference (RTPP)

Mice were placed in a custom-made plastic rectangular box (50 cm X 30 cm) with no distinguishing cues. One side of the chamber was randomly designated to be the side where the lasers would be activated. Individual mice were initially placed in the chamber and allowed to explore before beginning the experiment. During the 30-minute session the laser would be activated (1Hz: 0.5 s pulse duration, light intensity between 15–20 mW) each time the mouse entered the laser-paired side of the chamber, and would be turned off when the mouse crossed back into the non-stimulation side of the chamber. The difference score was calculated by taking the time the mouse spent in the Laser side minus time in the No Laser side.

Mice expressing ChR2 or GFP in astrocytes were connected to a 473 nm laser using a 1×1 rotary joint. For mice expressing ChR2 in astrocytes and NpHR3.0 in GABA cells we used a 2×1 rotary joint connected to 405 nm and 593 nm laser. This allowed us to activate either ChR2 or NpHR3.0 individually or together. The same set up was used for all RTPP experiments.

#### Laser Conditioned Place Avoidance (Laser CPA)

Laser CPA experiments were performed over a four-day period using a two-chamber conditioning apparatus (Med Associates) with each chamber consisting of a black chamber with a vertical metal bar floor, and a white chamber with a mesh floor. One mouse with a strong initial preference for either chamber (defined as a mouse spending more than 75% of the time in either chamber) was discarded from analysis. On day 1 (pre-test), mice were placed in one of the two sides and allowed to freely explore the entire apparatus for 30 minutes. During days 2 and 3 (conditioning) mice were confined to one of the two chambers (counterbalanced across all mice) and received optical stimulation (0.5 seconds pulse, 18 mW) once a minute for 30 minutes, or no stimulation. During the no stimulation session mice were connected the same as during the laser session, but the laser was off. A minimum of 4 hours after the first training session mice were placed into the other side of the chamber and the treatment reversed. Day 4 (posttest), mice were allowed to freely roam the apparatus for 30 minutes. The difference score was calculated by taking the time each mouse spent on the laser paired side in the post-test minus the time spent in the pre-test.

#### Cocaine Conditioned Place Preference (cocaine CPP)

A modified version of Laser CPA and the same chamber were used to perform cocaine CPP experiments. The pre-test and the post-test were performed identically, but the conditioning sessions were changed. During days 2 and 3 of conditioning, mice were confined to one of the two chambers (counterbalanced across all mice) and received an i.p. cocaine (15 mg/kg) injection paired with optical stimulation (0.5 seconds pulse, 18 mW) once a minute for 30 minutes, or i.p. saline injection with no laser stimulation. During both conditioning sessions mice were tethered to the patch cable. The difference score was calculated by taking the time each mouse spent on the cocaine/laser paired side in the post-test minus the time spent in the pre-test.

#### Brain slice preparation and electrophysiology

After completion of behavioral experiments, adult 10 to 14 week old mice were deeply anesthetized with isoflurane followed by rapid decapitation. The brain was quickly removed and chilled in ice cold cutting solution containing (in mM): 110 cholineCl, 2.5 KCl, 1.25 NaH_2_ PO_4_, 7 MgCl_2_, 0.5 CaCl_2_, 10 dextrose, 25 NaHCO_3_, and oxygenated with 95% O_2_ −5%CO_2_. 250 μm horizontal brain slices containing VTA were incubated for 30 minutes at 35° C in a chamber filled with artificial cerebrospinal fluid (aCSF) and stored at room temperature for the remainder of the day. Slices were transferred to a recording chamber where they continuously received aCSF (33–35° C) at a rate of 2 ml per minute by a gravity-feed system. aCSF was composed of (in mM): 126 NaCl, 2.5 KCl, 1.25 NaH_2_ PO_4_, 2 MgCl_2_, 2 CaCl_2_, 10 dextrose, 25 NaHCO_3_, was equilibrated using 95% O_2_ −5%CO_2_ and had a pH of 7.2.

Midbrain neurons and astrocytes were visualized using gradient contrast illumination through a 40X water-immersion lens attached to an Olympus BX51 upright microscope. The VTA was defined as neurons medial to the medial terminal nucleus of the accessory optic tract. Neurons were considered dopaminergic based on the following electrophysiological criteria: have a hyperpolarization-activated inward current, slow spontaneous firing frequency ≤ 5 Hz and an action potential half-width ≥ 1.5 ms. Dopamine (TH-positive) and GABA (GABA-positive) neurons filled with biocytin were further identified immunocytochemically (Fig. S4). Astrocytes expressed GFP and were visualized with a 40X or 60X water-immersion objective on a fluorescent microscope capable of detecting GFP.

Whole cell recording pipettes with a tip resistance between 4–6 MOhm were pulled from borosilicate glass using a P-97 Flaming/Brown electrode puller (Sutter Instruments). For voltage-clamp experiments recording pipettes were filled with internal solution containing (in mm): 110 d-gluconic acid, 110 CsOH, 10 CsCl_2_, 1 EGTA, 10 HEPES, 1 ATP, 10 mm phosphocreatine. Pipettes used for current clamp recordings contained (in mM): 138 K-gluconate, 10 HEPES, 2 MgCl_2_, 0.2 EGTA, 0.0001 CaCl_2_, 4 Na-ATP, 0.4 Na-GTP.

Postsynaptic currents (PSC) from excitatory and inhibitory afferents were evoked with no blockers present by placing a 250 μm bipolar stimulating electrode within 500 μm of the neuron being recorded and applying a single pulse (0.05 ms). PSCs were isolated by voltage clamping the cells at −55 mV for EPSCs and at +10 mV for IPSCs. The effect astrocytes had on synaptic activity was tested by photoactivating ChR2 on astrocytes for 500 ms (constant stimulation) and evoking PSCs 400 ms into the optical stimulation period. ChR2 on astrocytes was stimulated using a 473 or 405 nm laser (10–20 mW). Afferent-mediated currents on astrocytes were obtained by applying 50 Hz electrical stimulation (20 pulses, 0.05 ms pulse width). Whole cell recordings were made using a Multiclamp 700B amplifier (Molecular Devices). Signals were digitized at 50 kHz and saved to a hard drive for analysis using Axograph X (Axograph).

#### Statistics

A student’s two-tailed t-test was used when comparing between two groups. A One-way ANOVA with Tukey’s post hoc test was used for comparisons across multiple groups. A P-value < 0.05 was used as our significance level for all tests. Power analysis showed greater than 70 % power in all analyses.

## Data availability

The data supporting these findings are available from the corresponding author upon request.

## SUPPLEMENTARY FIGURES

**Figure S1.**
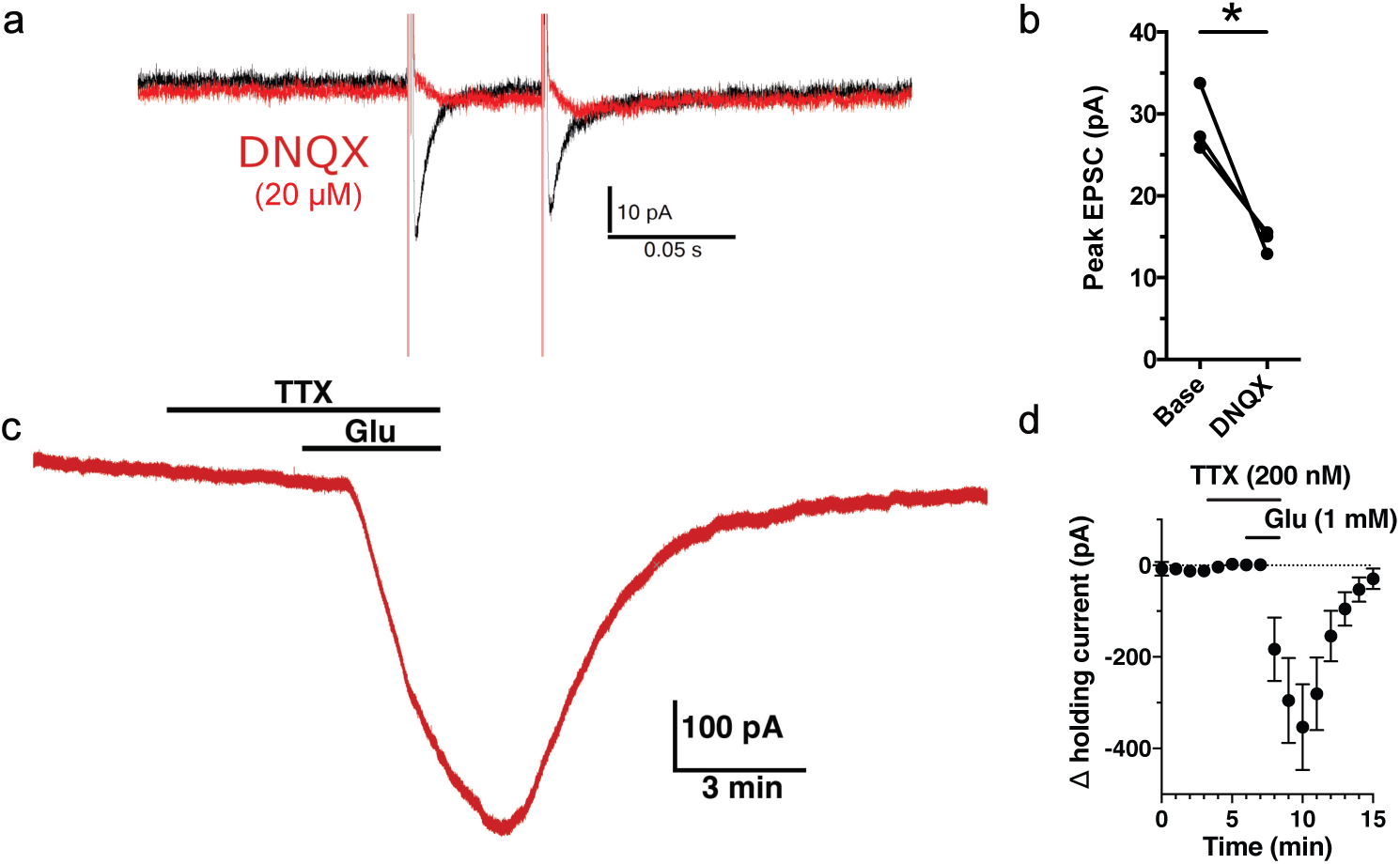
VTA Astrocytes are sensitive to glutamate via AMPA receptor activation. **(a)** Electrical stimulation during a whole-cell voltage clamp astrocyte recording evokes fast inward current responses (black traces). Bath application of the AMPA receptor blocker, DNQX (20 μM), blocks the response to electrical stimulation (red trace). **(b)** Summary data (paired t-test t_(2)_ = 4.473, P = 0.047, n = 3 cells from 2 mice). **(c)** Bath application of glutamate (1 mM) in the presence of TTX (200 nM) during a whole-cell recording of an astrocyte elicits a large inward current, indicating the presence of glutamate receptors on the recorded astrocyte. **(d)** Summary data (Repeated Measures t_(6)_ = 4.700, P = 0.003, n = 7 cells from 3 mice).

**Figure S2.**
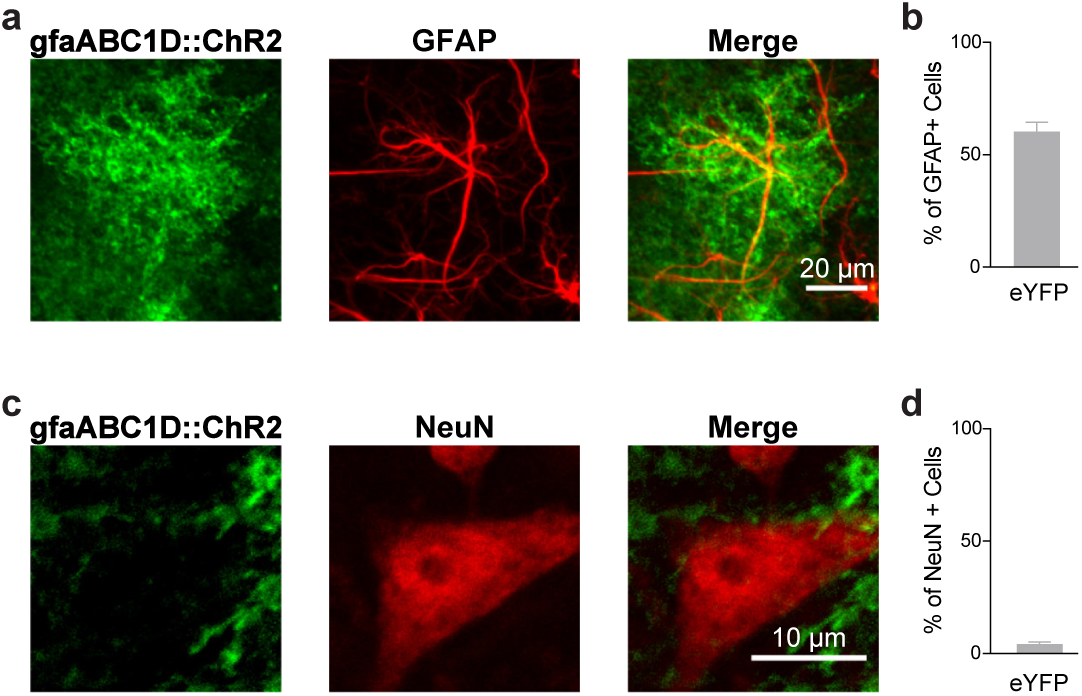
Channelrhodopsin is specifically expressed in astrocytes, not in neurons. **(a)** Immunocytochemistry image showing a cell expressing both gfaABC1D-eYFP (green) and GFAP (red). On the right are the two images merged together. **(b)** Quantification of the percentage of GFAP positive cells expressing eYFP (n = 4 mice). **(c)** Image showing a NeuN positive cell (red) surrounded by eYFP (green). **(d)** Quantification of the percentage of NeuN positive cells expressing eYFP (n = 3 mice).

**Figure S3.**
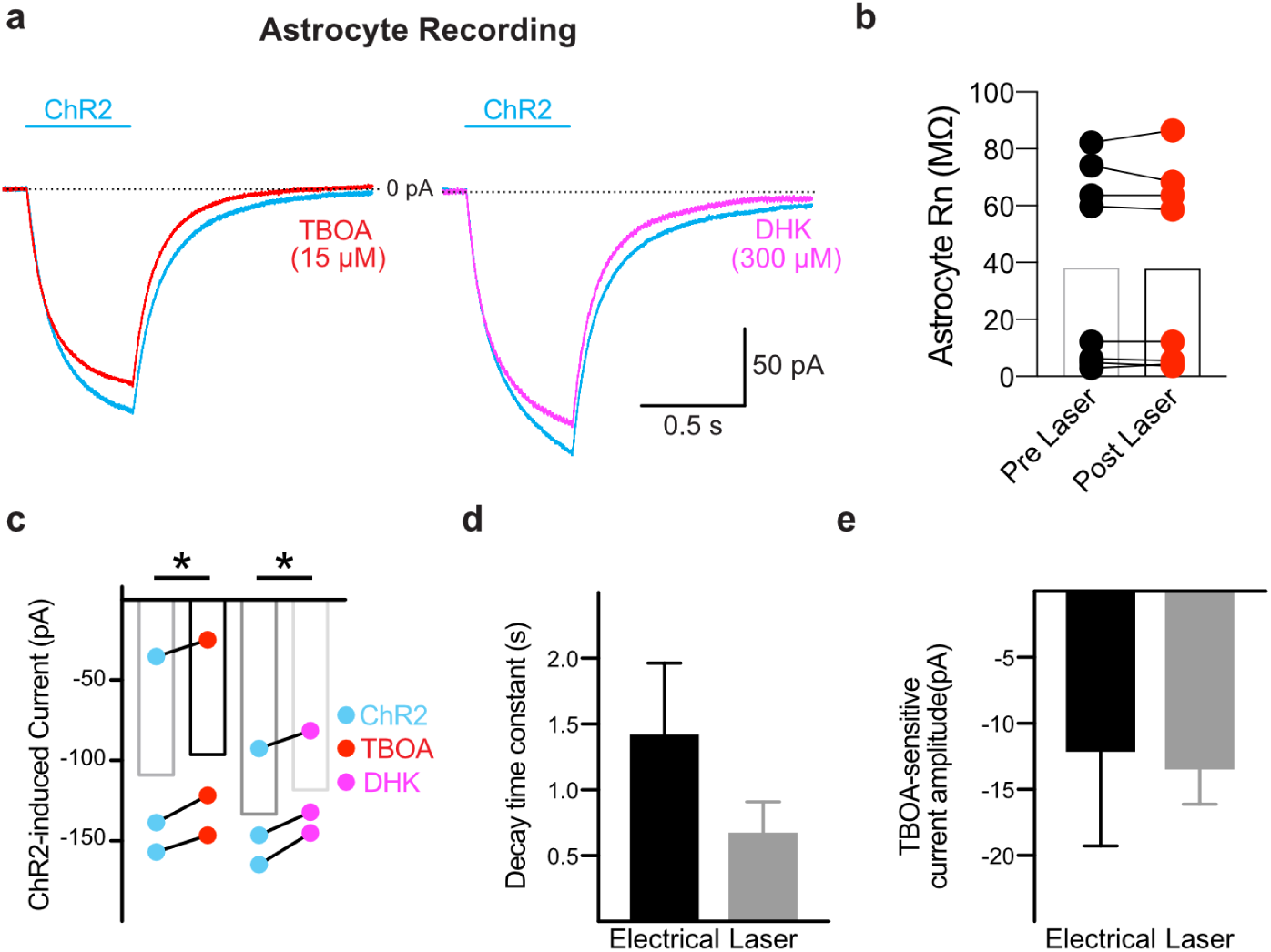
ChR2-mediated currents in VTA astrocytes possess properties similar to electrical stimulation. **(a)** Whole-cell recording from a VTA astrocyte showing a ChR2-mediated current (blue; no laser = 0.8325 ± 0.6587 pA, ChR2 = −87.06 ± 19.56 pA; paired t-test t_(8)_ = 4.437, P = 0.002, n = 9 cells, 4 mice; summary not shown) and the current remaining after application of 15 μm TFB-TBOA (left red) or 300 μm DHK (right magenta). **(b)** Summary of the astrocyte input resistance before and after stimulation of ChR2 in VTA astrocytes, indicating astrocytes remain healthy following photoactivation. **(c)** Summary of the ChR2 current produced in VTA astrocytes before (blue) and after (red) addition of TFB-TBOA (paired t-test t_(2)_ = 5.607, P = 0.03, n = 3 cells) or after (magenta) addition of DHK (paired t-test t_(2)_ = 6.051, P = 0.02, n = 3 cells, 2 mice). **(d)** Decay time constant (Tau) for the TBOA-sensitive current elicited by electrical stimulation of the slice (black bar) and ChR2 laser stimulation (gray bar) of the recorded astrocyte (unpaired t-test t_(6)_ = 0.3568, P = 0.35, n = 3, n = 5 cells respectively, 2 mice). **(e)** Peak amplitudes of the TFB-TBOA-sensitive currents (unpaired t-test t_(6)_ = 0.1386, P = 0.89, n = 3 and n =5 cells respectively, 2 mice). **: P < 0.01, *: P < 0.05; Error bars indicate ± SEM.

**Figure S4.**
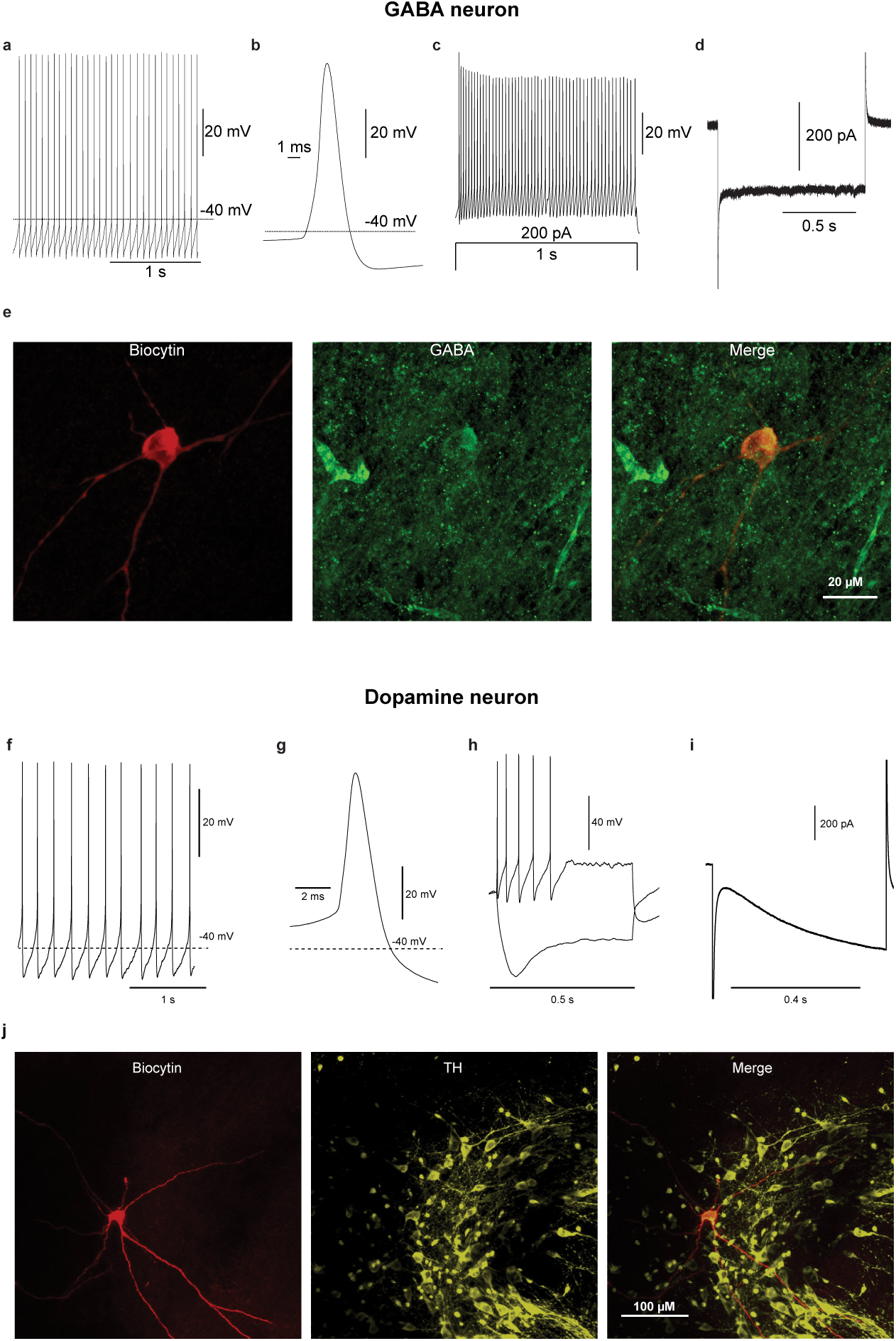
Identification of VTA neurons. Whole-cell current-clamp recording from a GABA (a-e) neuron and dopamine neuron (f-i). (**a, b**) One of the action potentials from (a) was expanded to emphasize the action potential half width (average action potential half width = 0.99 ms). (**c**) Recording showing no spike frequency adaptation in response to a 200 pA current injection. **(d)** Voltage clamp recording showing lack of Ih in response to hyperpolarizing the cell from −60 mV to −120 mV for 1 second (average Ih = −37.9 pA). (**e**) Immunohistochemistry images taken with a 40X objective showing a biocytin filled cell visualized using Alexa 594 (red) and GABA visualized using Alexa 488 (green). On the right is a merged image to show co-localization between the two fluorophores. **(f, g)** One of the action potentials from (f) was expanded to emphasize the wide action potential half width (average half width = 3.37 ms). **(h)** Example recording showing spike frequency adaptation in response to a 200 pA current injection and a characteristic sag in response to hyperpolarizing current injection (h; −200pA). **(i)** Developing Ih due to voltage clamp step (average Ih = −156.4 pA). **(e)** Immunohistochemistry images (20X objective) showing a biocytin-filled cell (red) with overlapping tyrosine hydroxylase staining (yellow). On the right is a merged image showing co-localization of the two fluorophores.

**Figure S5.**
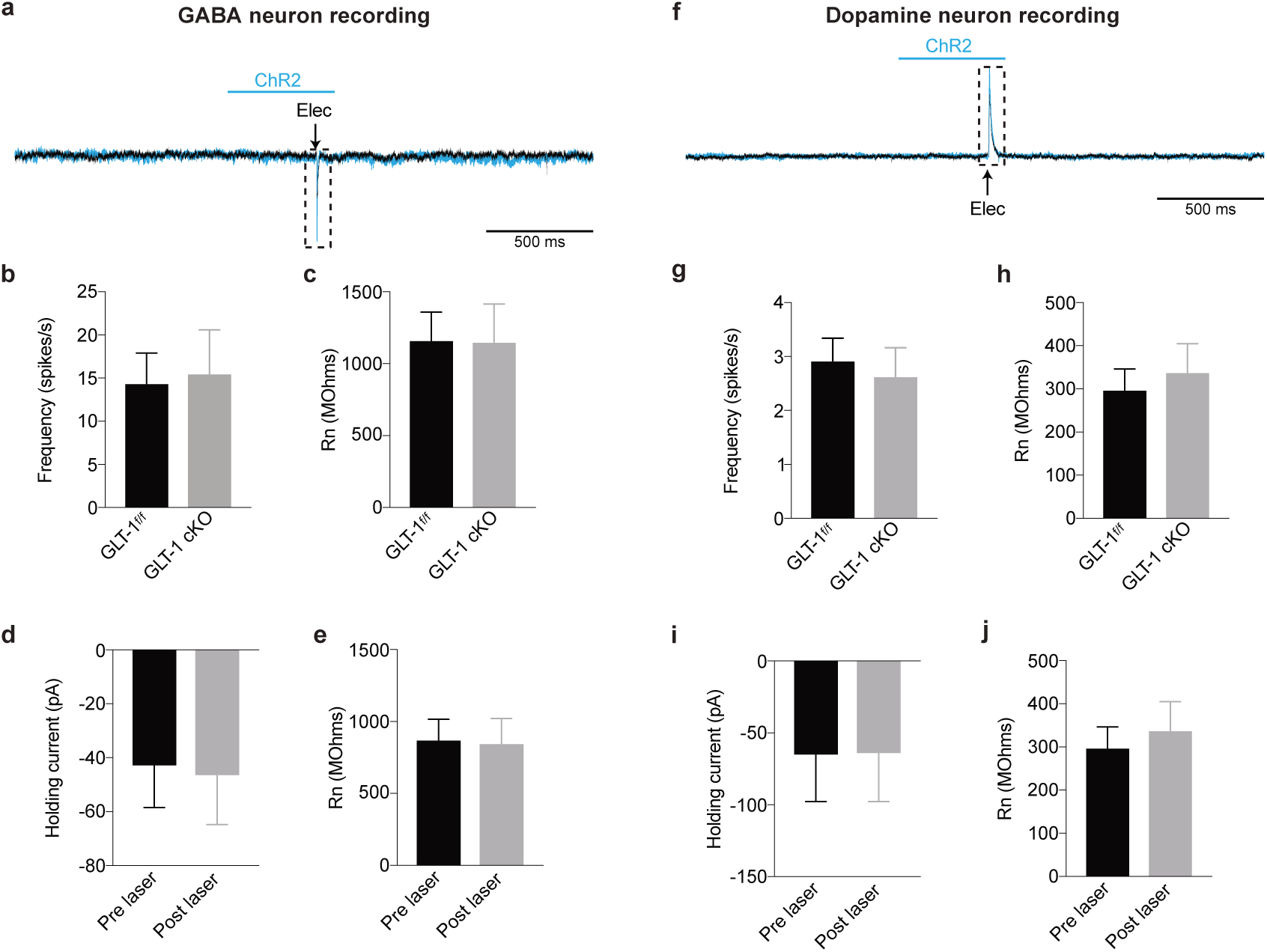
The intrinsic properties of VTA neurons are not altered by removing GLT-1 from astrocytes or by photoactivation of astrocytes. **(a)** Recording from a GABA neuron where astrocyte activation increases the EPSC. Summarized data showing there is no difference in **(b)** firing frequency (unpaired t-test t_(8)_ = 0.1831, P = 0.85, GLT-1^f/f^ n = 5 cells, 3 mice; GLT-1 cKO n = 5 cells, 2 mice) or **(c)** input resistance (unpaired t-test t(i2) = 0.0282, P = 0.97, GLT-1^f/f^ n = 6, 3 mice; GLT-1 cKO n = 8 cells, 2 mice) between GLT-1^f/f^ or GLT-1 cKO mice. Summarized data from GABA neuron showing there is no change in the **(d)** holding current (paired t-test t_(8)_ = 1.166, P = 0.27, n = 9 cells, 4 mice) or **(e)** input resistance (paired t-test t_(8)_ = 0.2121, P = 0.83, n = 9 cells, 4 mice) before and after astrocyte activation. **(f)** Recording from a dopamine neuron where astrocyte activation increases the IPSC. Summarized data from dopamine cells showing there is no difference in **(g)** firing frequency (unpaired t-test t_(11)_ = 0.4171, P = 0.68, GLT-1^f/f^ n = 6 cells, 2 mice; GLT-1 cKO n = 7 cells, 2 mice) or **(h)** input resistance (unpaired t-test t_(13)_ = 0.4818, P = 0.63, GLT-1^f/f^ n = 7 cells, 2 mice; GLT-1 cKO n = 8 cells, 2 mice) between GLT-1^f/f^ or GLT-1 cKO mice. Summarized data from dopamine neuron showing there is no change in the **(i)** holding current (paired t-test t_(5)_ = 0.5782, P = 0.58, n = 6 cells, 3 mice) or **(j)** input resistance (paired t-test t_(5)_ = 1.763, P = 0.13, n = 6 cells, 3 mice) before and after astrocyte activation.

**Figure S6.**
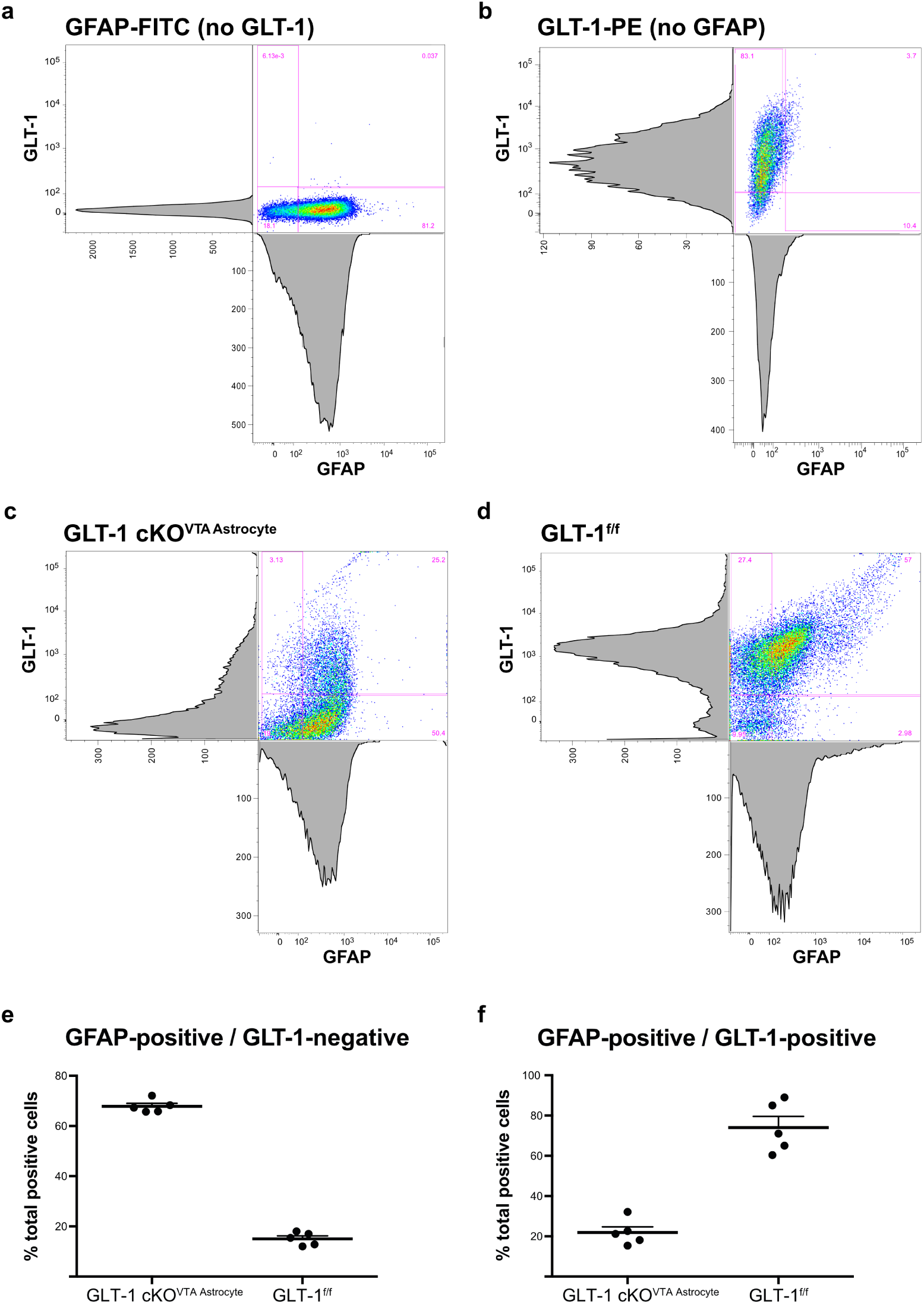
Loss of GLT-1 expression from VTA astrocytes. **(a-d)** Flow cytometry color dot-plots and histograms for GLT-1 (y-axis) and GFAP (x-axis) fluorescence intensities. **(a)** Control plot with no GLT-1 staining. **(b)** Control plot with no GFAP staining. **(c)** Flow cytometry from a GLT-1 cKOVTA Astrocyte (GLT-1^f/f^, gfaABC1D::Cre^VTA^) mouse with GLT-1 histogram peak indicating a shift toward GLT-1 negative cells relative to a GLT-1^f/f^ mouse (d). The histogram peak for GFAP is not different from the control histogram in (a), indicating the specificity of the knockout. **(d)** GLT-1 and GFAP histograms from a GLT-1^f/f^ mouse illustrates a large proportion of GLT-1 and GFAP positive cells (upper right quadrant). **(e, f)** Summary data indicating efficiency (e) and specificity (f) of the conditional knockout.

**Figure S7.**
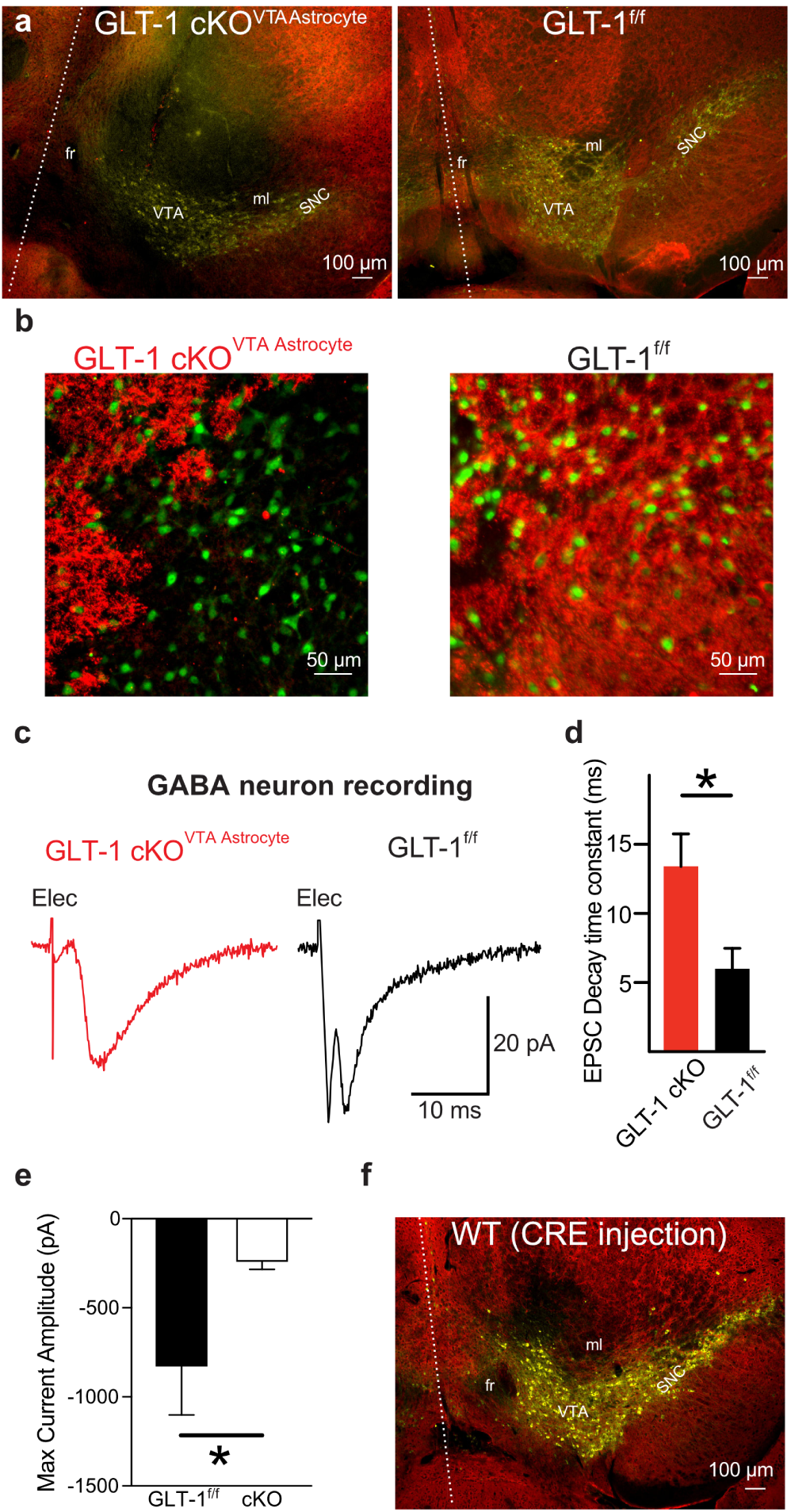
Loss of GLT-1 increases the decay kinetics and decreases maximum current generated on VTA GABA neurons. **(a)** Images (4X objective) illustrating loss of GLT-1 expression (red) in the VTA (GLT-1 cKO^VTA Astrocyte^, left) versus a mouse with a control injection (GLT-1^f/f^, right). For both (a) and (b) yellow is TH expression. **(b)** Image (20 X) showing NeuN positive cells (green) with GLT-1 staining (red). **(c)** Recordings from GABA neuron EPSC’s in a GLT-1 cKO^VTA Astrocyte^ (red; edge of injection site) and a GLT-f (black) mouse. **(d)** GLT-1 cKO^VTA Astrocyte^ mice (red trace) have a slower decay time constant of the EPSCs compared to GLT-1^f/f^ mice (black trace; unpaired t-test t_(9)_ = 2.531, P = 0.03, n = 6 and n = 5 cells respectively, 2 mice each). **(e)** Summarized data showing a difference in the maximum current generated when 200 μM glutamate is uncaged onto GABA neurons recorded from GLT-1^f/f^ and GLT-1 cKO mice (unpaired t-test t_(15)_ = 2.548, P = 0.022, n = 10 and n = 7 cells respectively, 3 mice each). **(f)** 4X image illustrating normal GLT-1 expression (red) when Cre is driven in a wild type (WT) mouse (GLT-1^+/+^, gfaABC1D::Cre^VTA^). fr = fasciculus retroflexus; ml = medial lemniscus; SNC = substantia nigra pars compacta; VTA = ventral tegmental area. Dashed line indicates midline. All images dorsal is up. Error bars indicate ± SEM.

**Figure S8.**
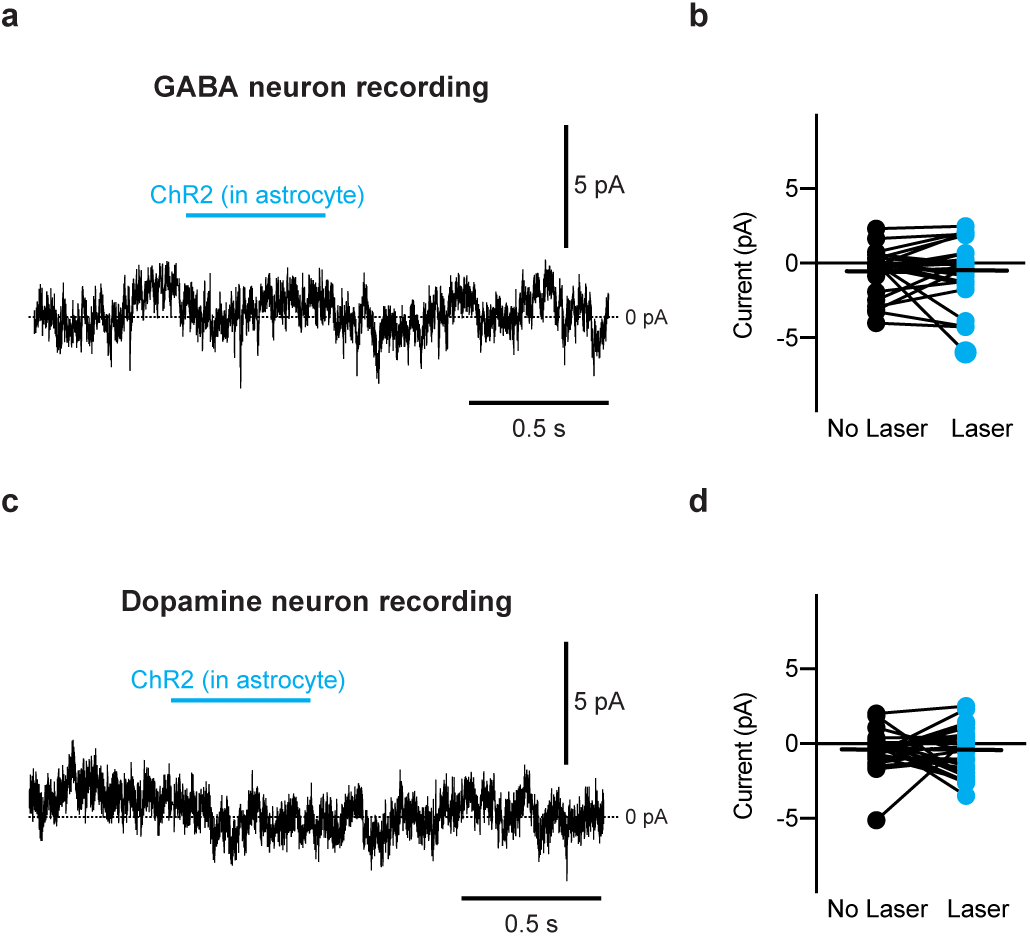
Astrocyte activation does not directly affect GABA or dopamine neurons. (**a**) Recording from a VTA GABA and **(c)** dopamine neuron while activating astrocytes expressing ChR2 (500 ms constant light pulse). Summary data comparing the average holding current 100 ms before astrocyte stimulation with the last 100 ms of astrocyte stimulation for recordings from **(b)** GABA (paired t-test t_(24)_ = 0.215, P = 0.98, n = 25 cells, 9 mice) and **(d)** dopamine neurons (paired t-test t_(28)_ = 0.2414, P = 0.811, n = 25 cells, 11 mice).

**Figure S9.**
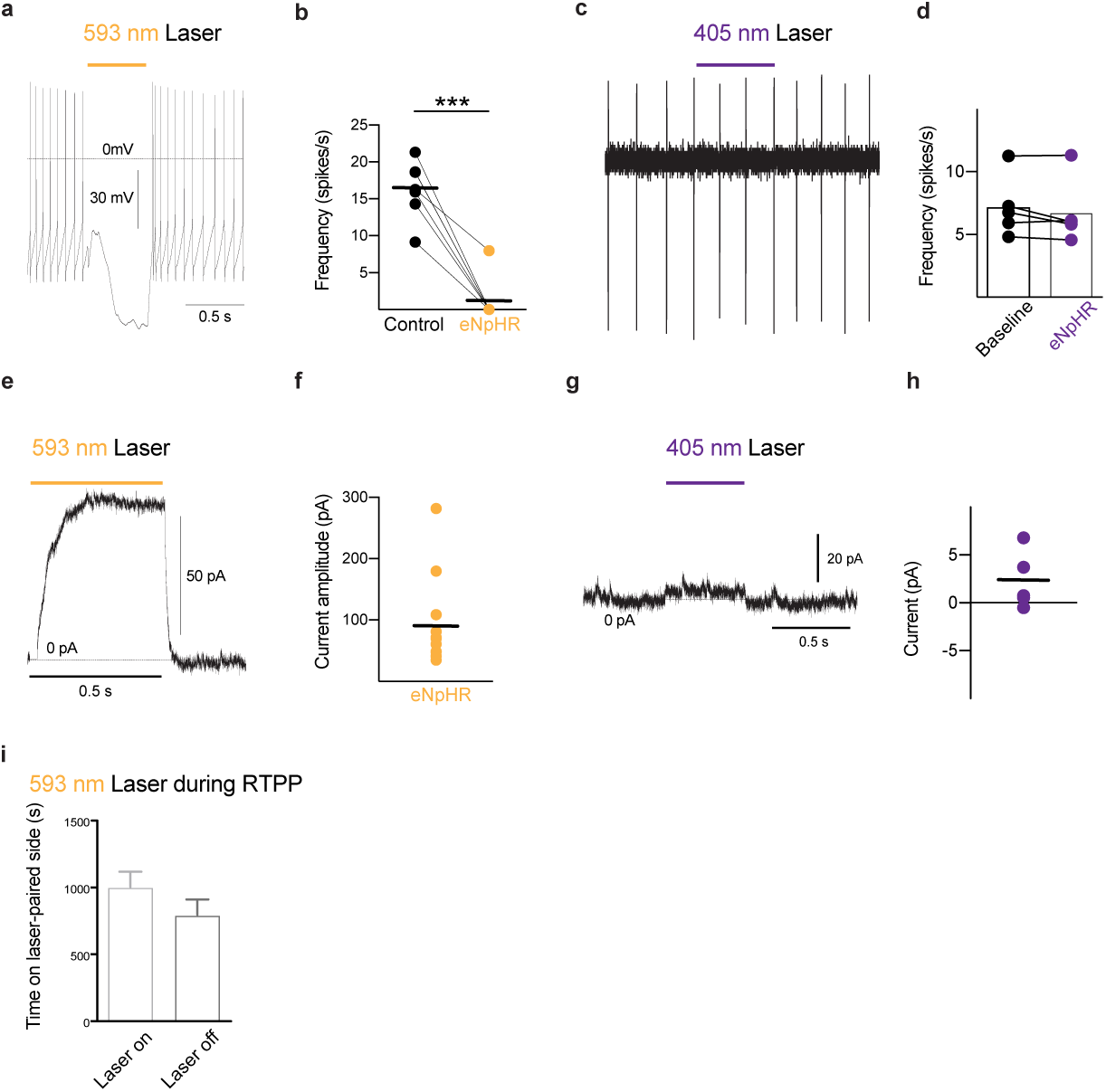
Photoactivation of eNpHR3.0 by 593 nm laser, but not a 405 nm laser, inhibits firing of GABA neurons. **(a)** Whole cell current-clamp recording from a VTA GABA neuron obtained from a Vgat-Cre mouse expressing eNpHR3.0 in GABA neurons. Photoactivation of eNpHR3.0 using a 593 nM laser prevents firing of action potentials in GABA neurons. **(b)** Summarized data showing the effect eNpHR3.0 (yellow) has on the firing of action potentials in GABA neurons (paired t-test t_(6)_ = 8.141, P = 0.0002, n =7 cells, 3 mice). **(c)** Cell-attached recording from a GABA neuron where 405 nm laser was applied with **(d)** accompanying summarized data (paired t-test t_(4)_ = 1.562, P = 0.19, n = 5 cells, 2 mice). **(e)** Whole-cell voltage clamp recording from a GABA neuron where activation of eNpHR3.0 with a 593 nM laser produces an outward current with **(f)** a summary of the currents generated (paired t-test t_(10)_ = 4.01, P = 0.002, n =11 cells, 3 mice). **(g)** A voltage-clamp recording from the same GABA neuron as (c) where activation of a 405 nm laser with equivalent light intensity does not change the firing frequency of GABA neurons. **(h)** Summary of the outward current produced by a 405 nm laser on GABA neurons (paired t-test t_(4)_ = 1.523, P = 0.20, n = 5 cells, 2 mice). **(i)** Summarized data demonstrating GABA neuron hyperpolarization does not produce rewarding affects in the RTPP behavior assay (paired t-test t_(6)_ = 0.9036, P = 0.4, n = 7 mice). ***: P < 0.001

**Figure S10.**
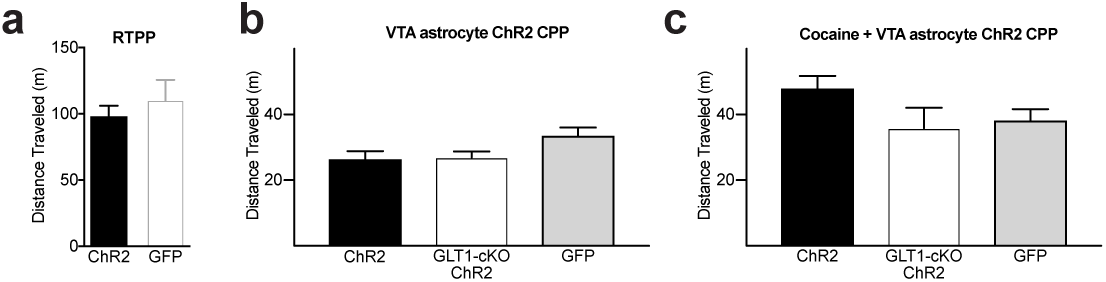
Astrocyte activation does not alter locomotor activity in real-time place preference or conditioned place preference assays. **(a)** There was no difference in the total distance traveled between mice expressing GFP or ChR2 in VTA astrocytes during a 30 minute RTPP session (unpaired t-test t_(15)_ = 0.7177, P = 0.48, n = 8 and n = 9 mice respectively). (b) Comparing the total distance traveled between mice expressing either ChR2, GLT-1 cKO with ChR2, or GFP in VTA astrocytes during laser activation (One-way ANOVA, F_(2,33)_ = 2.203, P = 0.1264). (c) Comparing the total distance traveled between mice expressing either ChR2, GLT-1 cKO with ChR2, or GFP in VTA astrocytes during cocaine + laser activation (One-way ANOVA, F_(2,27)_ = 2.299, P = 0.1197); Error bars indicate ± SEM.

**Supplementary Video 1**.

**ChR2 stimulation on VTA astrocytes elicits avoidance**. Mouse receives ChR2 photoactivation of VTA astrocytes (1Hz: 0.5 s pulse duration, 15–20 mW) when venturing to the top half of the field. Movie (4X speed) is 15 s of the 30 minute session.

## ACKNOWLEDGEMENTS

We thank J. Wadiche, C. Wilson and M. Higgs for helpful comments. Funding: This research was supported by National Institutes of Health Grants MH079276 (CAP), MH113341 (CAP), DA030530 (CAP), DA033386 (MJW), DA042362(MJW), GM060655 (JAG), F31DA041303 (JAG), NS066019 (PAR), MH104318 (PAR), DA032701 (MJB) and National Science Foundation Grant HRD-1249284 (JAG). Imaging data collection was supported by Grant G12MD007591.

## Contributions

JAG, JMP, GMB, KT, AMM, MJB, CWT and EAS collected data. JAG, EAS, KT, AMM, JMP, MJW, MJB, CWT and CAP analyzed data. JAG, JMP, GMB, EAS, KT, MJB, CWT, NBC, and PAR prepared animals and reagents. JAG, SAQ, and CAP wrote the paper.

## Competing interests

The authors declare no competing interests

